# Design of an imaging probe to monitor real-time redistribution of L-type voltage gated calcium channels in astrocytic glutamate signalling

**DOI:** 10.1101/2020.11.19.390013

**Authors:** Mitra Sadat Tabatabaee, Jeff Kerkovius, Frederic Menard

## Abstract

**Purpose:** In the brain, astrocytes are non-excitable cells that undergo rapid morphological changes when stimulated by the excitatory neurotransmitter glutamate. We developed a chemical probe to monitor how glutamate affects the density and distribution of astrocytic L-type voltage-gated calcium channels (LTCC).

**Procedures:** The imaging probe FluoBar1 was created from a barbiturate ligand modified with a fluorescent coumarin moiety. The probe selectivity was examined with colocalization analyses of confocal fluorescence imaging in U118-MG and transfected COS-7 cells. Living cells treated with 50 nM FluoBar1 were imaged in real time to reveal changes in density and distribution of astrocytic LTCCs upon exposure to glutamate.

**Results:** FluoBar1 was synthesized in ten steps. The selectivity of the probe was demonstrated with immunoblotting and confocal imaging of immunostained cells expressing the Ca_V_1.2 isoform of LTCCs proteins. Applying FluoBar1 to astrocyte model cells U118-MG allowed us to measure a 5-fold increase in fluorescence density of LTCCs upon glutamate exposure.

**Conclusions:** Imaging probe FluoBar1 allows the real-time monitoring of LTCCs in living cells, revealing for first time that glutamate causes a rapid increase of LTCC membranar density in astrocyte model cells. FluoBar1 may help tackle previously intractable questions about LTCC dynamics in cellular events.

## INTRODUCTION

L-type voltage-gated calcium channels (LTCC) are membrane proteins that mediate calcium fluxes upon depolarization of a cell [1, 2]. While LTCCs were thought to be exclusive to electrically active cells such as neurons, they are now known to be also expressed in non-excitable cells with a range of functions beyond electrical activity [3]. For instance, they are involved in stem cell regulation [4], cell differentiation [5], transcription regulation [6], extracellular signaling [7], synaptogenesis [8, 9], cell migration, and neural myelination [10].

The physiological role of L-type voltage-gated calcium channels correlates tightly with their distribution at the cell membrane [1]. Pravettoni *et al.* pioneered investigations on the spatiotemporal dynamics of LTCCs in neuronal development [11], and their essential role has been reviewed recently [12, 13]. While valuable knowledge was gained about LTCCs’ translocation and distribution in development, the lack of tools to monitor these channels in real time has hampered their study in cellular events.

At a synapse, glutamate is the main excitatory neurotransmitter that transduces signals between neurons [14]. Astrocytes were long thought to be passive support cells in the brain, but it is now well established that glutamate initiates an intracellular calcium rise that leads to a rapid extension of their processes [15–17]. Yet, three decades after introducing the tripartite synapse model (where an astrocyte makes a direct contact with the pre-and post-synaptic neurons) [18], the molecular mechanisms of astrocytic glutamate signaling are still under investigation [19–21]. Following the first report by Cornel-Bell *et al.* showing that glutamate initiates a calcium-dependent response of cultured astrocytes [15, 16], several groups have examined how astrocytes respond to neuronal activity and neurotransmitter release *in vivo* [22–29]. In line with the mounting evidence that calcium-permeable or calcium-releasing glutamate receptors may be involved [19, 27, 29], we recently found that LTCCs participate to the glutamate-induced response in a model astrocyte cell U118-MG [26]. While we suspected that a rapid LTCC redistribution might be associated, the tools available could not answer our questions satisfactorily, and urged us to create an imaging probe that could monitor the channels’ exofacial distribution in real time.

The conventional technique to visualize a protein in a cell is immunocytochemistry (ICC), where a detectable antibody binds to an epitope of the target protein [30]. However, ICC can be prone to reproducibility issues due to non-selective binding of antibodies [19, 31–33]. In addition, if the epitope for the antibody is intracellular, the cells must be fixed and permeabilized, which *de facto* prevents investigating protein dynamics [34]. Recombinant fusion with a fluorescent protein (e.g., GFP) is a popular alternative technique for live cell imaging, but protocols to express a FP-fused LTCC are complex, time-consuming, and can perturb a protein’s function due to misfolding, incomplete post-translational modification, or improper transport [35]. Therefore, we aimed to create a stable, efficient and user-friendly fluorescent chemical imaging tool that targets LTCCs in living cells.

Our chemical imaging probe (FluoBar1) is based on the barbiturate class of pyrimidinetriones reported by Silverman *et al.* (**1**, Fig. 1) [36, 37]. The labeling process in cells is operationally simple and requires only a three-minute incubation time with 50 nM of FluoBar1. This probe allows us to monitor and quantify LTCC localization, density and distribution in real time. All studies were performed with the voltage-gated calcium ion channel isoform 1.2 of LTCC (Ca_V_1.2) as it is the most prevalent in astrocytes [47, 52]. Herein, we report the synthesis, validation, and application of probe FluoBar1 to measure time-dependent changes on membrane-bound LTCCs upon glutamate stimulation.

**Figure 1.**
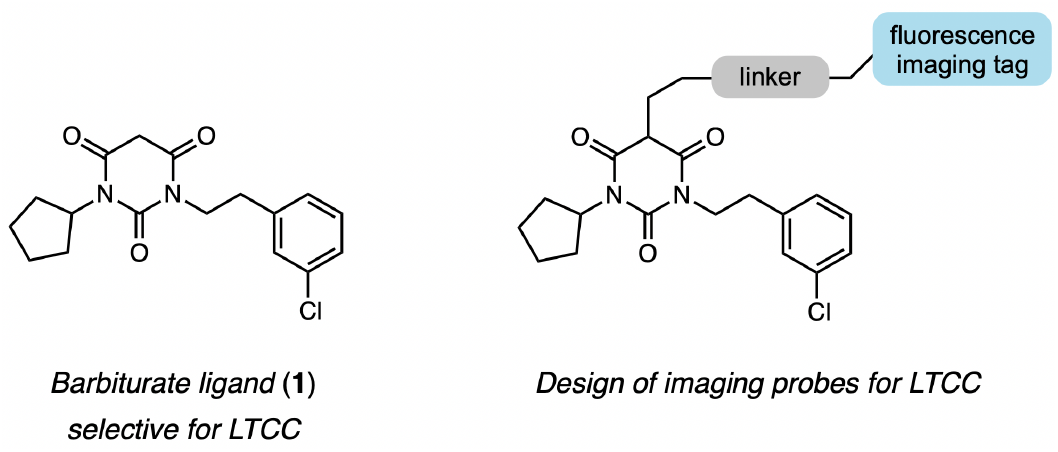
General structure of a fluorescence imaging probe based on the barbiturate class of pyrimidinetriones.

## MATERIAL AND METHODS

### Chemical synthesis

An overview of the synthesis is presented in Figure 2. The convergent synthesis consists of ten reactions with six steps for the longest linear sequence (14% overall yield). Starting materials are: cyclopentane carboxylic acid (**2**), 2-(3-chlorophenyl) acetonitrile, and 2-aminoethanol. All procedures for the synthesis of the imaging probes, reagents, instruments, and chemical characterization are detailed in the accompanying *Supporting Information* file (Fig. S2). Only the last step is described below for space considerations.

**Figure 2.**
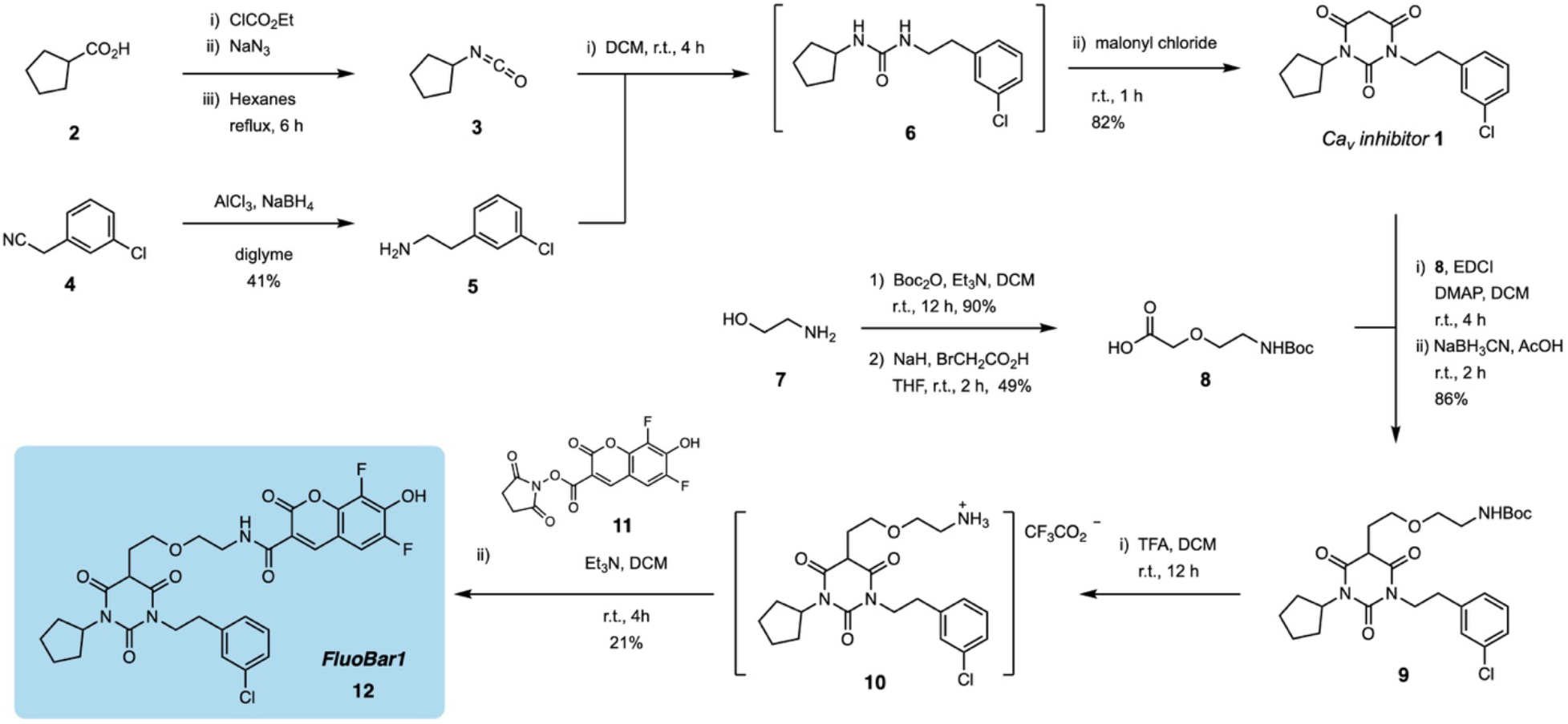
Synthesis of imaging probe FluoBarb1 (**12**). The fluorescent moiety is introduced at the last step to allow maximal flexibility in the type of desired reporter.

### Synthesis of FluoBar1 (12)

To a 250 mL flask was diluted the tethered barbiturate **9** (0.386 g, 0.741 mmol) in CH2Cl2 (40 mL). Trifluoroacetic acid (1.2 mL, 15.67 mmol) was added to the solution at r.t. and the reaction was stirred overnight. The next day, the flask was protected from light, triethylamine (4.2 mL, 30.1 mmol) was added, followed immediately by a solution of the NHS ester of Pacific Blue **11** (0.261 g, 0.765 mmol) in DMF (15 mL). The reaction was stirred for 4 hours at r.t. The reaction was quenched by slow addition into water (50 mL), followed by 1 M HCl aq. solution until the solution’s pH was less than 4. The mixture was extracted with ethyl acetate (3 x 40 mL). The combined organic layers were washed with water (2 x 40 mL), brine (50 mL), dried over anhydrous MgSO4, filtered, and concentrated under reduced pressure. The crude brown oil was purified by flash chromatography on silica gel (70% EtOAc/hexanes). The yellow powder product was further purified by HPLC. A 50 mg/mL solution of the product was prepared in acetonitrile, then injected in 160 μL aliquots on an Agilent 1290 Infinity UPLC equipped with a Waters Nova-Pak® HR C18 6 μm 19×300 mm prep-HPLC column; and eluted with an isocratic mixture of 70% acetonitrile containing 0.1% formic acid and 30% water containing 0.2% formic acid at a rate of 3.0 mL/min. The FluoBar1 compound was detected at 365 nm on a diode array detector, and eluted at 35 minutes. The collected fractions were lyophilized in the dark to yield FluoBar1 (**12**) as a light yellow/green powder (93 mg, 21%). ^**1**^**H NMR** (400 MHz, CDCl_3_) δ = 8.84 (br s, 1H), 8.69 (d, *J* = 1.4 Hz, 1H), 7.21-7.11 (m, 5H), 5.13 (p,*J* = 8.5 Hz, 1H), 4.06 (td, *J* = 7.8, 4.1 Hz, 2H), 3.59 (m, 7H), 2.85 (t, *J* = 7.4 Hz, 2H), 2.40 (m, 2H), 2.00-1.79 (m, 6H), 1.61-1.51 (m, 2H) ppm. ^**13**^**C NMR** (101 MHz, CDCl_3_) δ = 168.7, 168.5, 161.7, 160.1, 151.0, 148.7 (dd, *J* = 244.0, 3.6 Hz), 147.6, 140.8 (dd, *J* = 9.2, 1.7 Hz), 140.1, 139.6 (dd, *J* = 18.4, 12.8 Hz), 138.8 (dd, *J* = 249.7, 5.8 Hz), 134.4, 129.9, 129.2, 127.2, 126.7, 116.7, 110.5 (d, *J* = 9.4 Hz), 110.0 (dd, *J* = 20.2, 3.5 Hz), 69.2, 67.3, 54.7, 46.2, 42.9, 39.8, 33.8, 30.1, 28.9, 28.8, 25.8, 25.7 ppm. ^**19**^**F NMR** (377 MHz, CDCl_3_) δ = –136.2 (t, *J* = 9.2 Hz), –152.7 (d, *J* = 8.8 Hz) ppm. **IR** (FT-IR, solid state): 3345, 3060, 2948, 2866, 1720, 1676 cm^−1^. **HRMS** (ESI-TOF) *m/z*: [M–H]^−^calcd for C31H29ClF2N3O8 644.1616; found 644.1619.

### Cell culture

Human astrocytoma U118-MG cells (HTB-15, ATCC) and human kidney COS-7 cells (CRL-1651, ATCC) were cultured in Dulbecco’s modified essential medium (DMEM, Gibco 11995-065) supplemented with 10% v/v heat-inactivated fetal bovine serum (Gibco 12483-020) 1% v/v penicillin 10,000 U/ml and streptomycin 10,000 μg/ml (Gibco 15140-163). Cells were incubated in a humidified atmosphere containing 5% CO2 at 37 °C, and typically passaged at 80-95% confluence using a 0.25% trypsin-EDTA solution pre-warmed to 42 °C (Gibco 25200-072) for a maximum of 5 minutes.

For experiments, cells from a 10 cm culture dish at ~80% confluence were resuspended with a pre-warmed 0.25% trypsin-EDTA solution and transferred to a 35 mm glass-bottom culture dish (Mutsunami D35-14-1) in phenol red-free, high glucose DMEM supplemented with 25 mM HEPES (Gibco 21063-029). Enough cells were transferred to obtain ~50% confluence after adhesion.

### Transfection

COS-7 cells were co-transfected with ~1200 ng each of two plasmids: pcDNA3-α1c (rat Ca_V_1.2) generously gifted by Terry P. Snutch (UBC Vancouver), and pcDNA3-EGFP was a gift from Doug Golenbock (Addgene plasmid #13031) using a calcium phosphate precipitation protocol [38]. pcDNA3-EGFP is used as reporter to identify cells that uptook pcDNA3-α1c (Ca_V_1.2).

### Immunocytochemistry

Cells were stained with antibodies raised against the α1c subunit of Ca_V_1.2 (rat), imaged by Olympus FV1000 fluorescence confocal microscopy, and analyzed for co-localization of fluorescence signals. Briefly: plated cells were washed with phosphate buffer saline (PBS) solution, then a set of dishes were fixed in 4% paraformaldehyde in PBS solution for 20 min at room temperature without stimulus; another set were exposed to glutamate for 10 minutes, then fixed with the same procedure. The fixed cells were incubated in a blocking solution (10% BSA in PBS), followed by overnight incubation at 4 °C with the primary antibody anti-Ca_V_1.2 (rabbit anti-rat/human polyclonal, Alomone ACC-003) in 3% BSA in PBS (1:200 dilution). After three rinses with PBS, cells were treated with a 1:200 dilution of the secondary antibody goat anti-rabbit IgG (H+L) conjugated to Dylight 550 (Thermoscientific PA5-84541) at room tempertature on a rocking platform for 2 hours. Selectivity of the Ca_V_1.2 antibody and its isotype was controlled with blotting assays based on the differential expression of native LTCCs in U118-MG, COS7, and HEK cells (see supp. info. Fig S1).

### Protein labelling with FluoBar1

Live imaging with labelled Ca_V_1.2 proteins was performed as follows: live cells were incubated with 50 nM of FluoBar1 in PBS solution at room temperature for 2-5 minutes. Short incubation times minimize passive diffusion of the probe in the cells membrane. After a single wash with PBS solution, 2.0 ml Hank’s balanced salt solution (HBSS, Lonza 10547F) supplemented with 2 mM CaCl2 (Sigma Aldrich, C1016) was added as medium for imaging. Fixed cells were imaged using the same protocol.

### Cell treatment with excitatory neurotransmitter

On the confocal microscope stage, U118-MG cells were first stained with FluoBar1 as described above in HBSS supplemented with 2 mM CaCl2 (Sigma Aldrich C1016). Imaging began 10 minute before stimulating cells by adding a glutamate (Alfa Aesar A15031) aliquot to obtain a final dish concentration of 100 μM glutamate. More specifically, a 20x stock solution of L-glutamate was prepared in deionized millipore water and filter-sterilized through a 0.22 μm syringe filter [15, 17]. A 10.0 μL aliquot of the 20x glutamate solution was carefully delivered to the medium (2.0 mL) at the edge of the 35 mm dish. A concentrated stock solution was used to avoid osmotic shock.

### Confocal live imaging

Live U118-MG cells pretreated with FluoBar1 were imaged every 20 seconds for a total number of 60 frames on an Olympus FV1000 fluorescence confocal microscope equipped with a Plan-ApoN 60x/i.4 oil-immersion objective. Photobleaching was minimized by irradiating samples for 1 second at 20 second intervals with the laser power set at 3%. Fluorescence of the chemical probe was excited with a 405 nm He– Ne laser (Olympus filter set AlexaFluor 405).

### Colocalization test

Fixed cells immunostained for Ca_V_1.2 (secondary antibody conjugated to Dylight 550) were incubated with 50 nM of FluoBar1 in PBS at room temperature on the confocal microscope stage. The probe solution was replaced with PBS solution after 3 minutes, and cells were imaged immediately thereafter. Images were acquired sequentially using fluorescence filters in separate channels: AlexaFluor405 for the probe (emission at ~ 450 nm, blue) and Cy3 for antibody–DyLight 550 (emission at ~620 nm, red). The images were acquired at 60x magnification, with Kalman factor 4, and analyzed for colocalization.

### Image analysis

Confocal microscopy images from AlexaFluor405 and Cy3 channels were processed with FIJI using the colocalization analysis function to obtain the Pearson’s *r* value, the Costes *P* value, and the Menders coefficient. The changes in fluorescence density of FluoBar1 were also measured with FIJI [39]. Briefly: each cell’s contour was outlined manually, the total fluorescence signal for this area was integrated, and the fluorescence density was calculated. Background fluorescence was measured by selecting three cell-free areas (approximately half a cell’s size each, from the same image) and their signal was integrated and their density was averaged. The area conversion value for calculatations was 3.02 pixels/μm. The corrected total cell fluorescence density (CTCF) was calculated according to: CTCF = (integrated density of a cell) – (integrated density of background).

All data were processed and plotted using Prism 7.04 from GraphPad. Each value represents the averaged changes measured from three separate culture dish; the results are presented as the mean ± S.E.M. Significant differences among groups were determined using parametric paired two-tailed T-test; a *p* value < 0.05 was considered significant.

## RESULTS

### Synthesis of FluoBar1

Imaging compound FluoBar1 (**12**) was prepared as outlined in Figure 2 with a convergent synthesis comprising a total of ten steps (see supporting information for detailed procedures and characterization). This route enables the introduction of a fluorescent moiety at the last step. A late-stage addition is advantageous as fluorophores are often costly, chemically sensitive, or difficult to isolate.

The first three steps are based on the combinatorial synthesis reported by Silverman and coworkers [36]. Briefly, alkyl carboxylic acid **2** was converted to the corresponding isocyanate using a Curtius rearrangement and was trapped directly with 3-chlorophenethylamine **5** to obtain unsymmetrical urea **6**. The urea was directly reacted with freshly prepared malonyl chloride to give barbiturate **1**. The latent nucleophilic character of the barbiturate’s dicarbonyl was exploited to append a carboxylic acid linker chain using a one-pot acylation/reduction sequence. Acid **8** was first activated with EDCI and was reacted with pyrimidone **1** to afford the ketone sidechain [40]. The resulting ketone was unstable to isolation, and was therefore reduced *in situ*. Reduction with sodium cyanoborohydride in acetic acid was found to afford the best compromise in terms of yield and purity. Deprotection of the linker’s carbamate under standard conditions afforded the trifluoroacetate salt **10** cleanly. Terminal ammonium **10** was reacted directly with the freshly prepared *N*-hydroxysuccinimic ester of Pacific Blue **11** [41] to yield the desired coumaryl amide FluorBar1 (**12**). The chemical probe is slightly light and air-sensitive: it was observed to show slow degradation by NMR after 24 h when maitained in solution at room temperature. It is stable for months when stored at –20 °C under nitrogen or argon.

### FluoBar1 is selective for Ca_V_1.2

Confocal fluorescence imaging of fixed U118-MG cells showed that FluoBar1 colocalized with the anti-Ca_V_1.2 antibody (Fig. 3). Colocalization analyses between fluorescence signals observed in a blue (FluoBar1, Olympus AlexaFluor405 filter) and a red channel (anti-Ca_V_1.2/DyLight550, Olympus Cy3 filter) returned a Pearson’s *r* value of 0.87 and a Costes *P* value of 1.00 (a *P* value above 0.95 confirms the accuracy of Pearson’s coefficient) (Table 1) [42] [43]. The Mender coefficient indicated >99% colocalization for both the blue and the red channels; this coefficient is the ratio of pixels from a channel that colocalized over the total signal observed the same channel for a given cell area [44]. The analyses suggest that unselective labeling by FluoBar1 is negligeable, or similar to the Ca_V_1.2 antibody used.

**Table 1.**
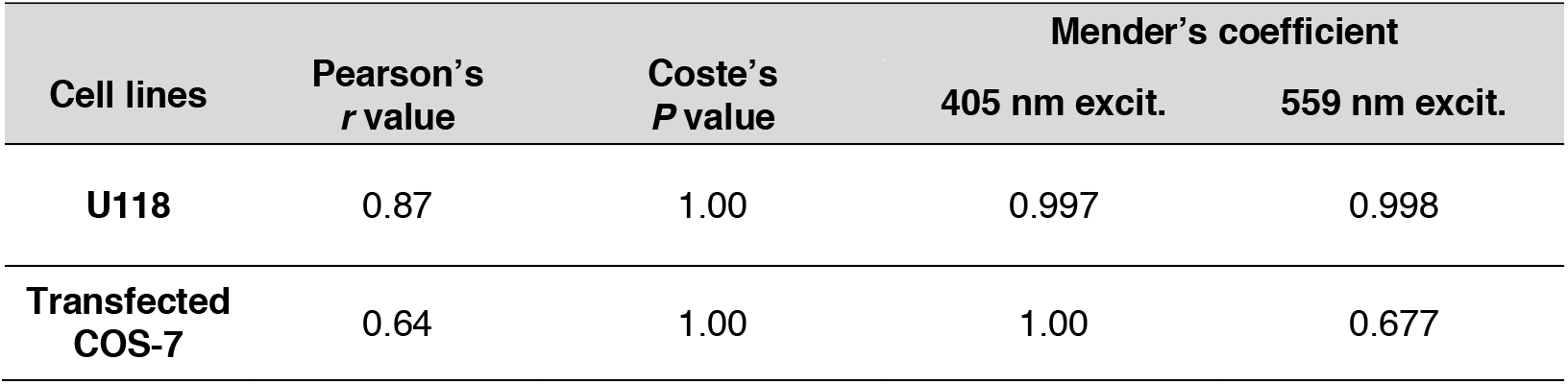
Colocalization analysis of confocal images. The fluorophores of FluoBarb1 and anti-Ca_V_1.2/DyLight550 colocalize with one another with high correlation.

| Cell lines | Pearson's *r* value | Coste's *P* value | Mender's coefficient |  |
| --- | --- | --- | --- | --- |
|  |  |  | 405 nm excit. | 559 nm excit. |
| **U118** | 0.87 | 1.00 | 0.997 | 0.998 |
| **Transfected COS-7** | 0.64 | 1.00 | 1.00 | 0.677 |

**Figure 3.**
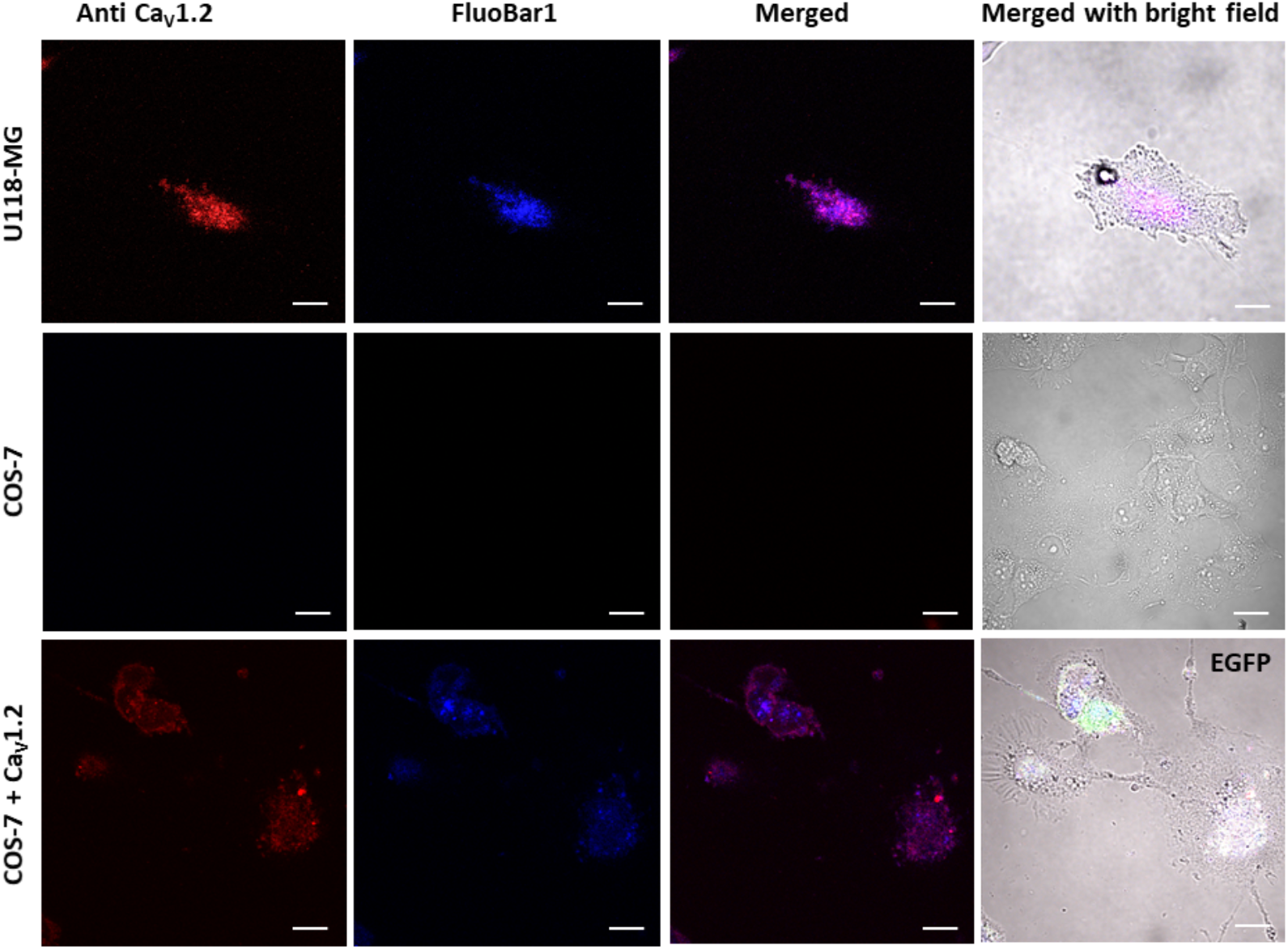
Representative confocal fluorescence images of cells labeled with an anti-Ca_V_1.2/DyLight550 antibody and FluoBar1. Imaging probe FluoBar1 is highly colocalized with the Ca_V_1.2 antibody in astrocytic U118-MG that natively express LTCCs, as well as in COS-7 cells transfected with Ca_V_1.2. Untransfected COS-7 do not express LTCCs and showed no significant labeling with either the probe or the antibody. Immunostained fixed cells were treated with 50 nM FluoBar1 for 3 minutes, then imaged in HBSS; 60x magnification, scale bars = 10 μm.

Negative controls with COS-7 cells showed no fluorescence signal above background with neither anti-Ca_V_1.2 nor FluoBar1. When transfected with a plasmid for the Ca_V_1.2 pore-forming subunit (α1C), COS-7 cells showed fluorescence signals with both FluoBar1 and the Ca_V_1.2/DyLight550 antibody (Fig. 3). The level of colocalization was lower than for the astrocyte model cells: a Pearson *r* value of 0.64 was calculated with high confidence (Costes *P* = 1.00).

### The distribution of LTCC changes when U118-MG cells respond to glutamate

FluoBar1 was applied to live astrocytic U118-MG cells to examine the possibility of monitoring Ca_V_1.2 proteins in real time (Fig 4). The labelled cells were exposed to 100 μM glutamate and time-lapse confocal imaging revealed a gradual increase in fluorescence density that leveled to a plateau after 10 minutes (from 5,299 ± 1,475 to 24,400 ± 6,482 AU/μm^2^, n = 3, Fig. 4A and 4B). Unpaired two tail *T*-test analysis shows a statistical significance P value < 0.05 (Fig. 4B). Along with fluorescence density increase, the patterns of Ca_V_1.2 distribution were observed to change rapidly upon exposure to glutamate.

**Figure 4.**
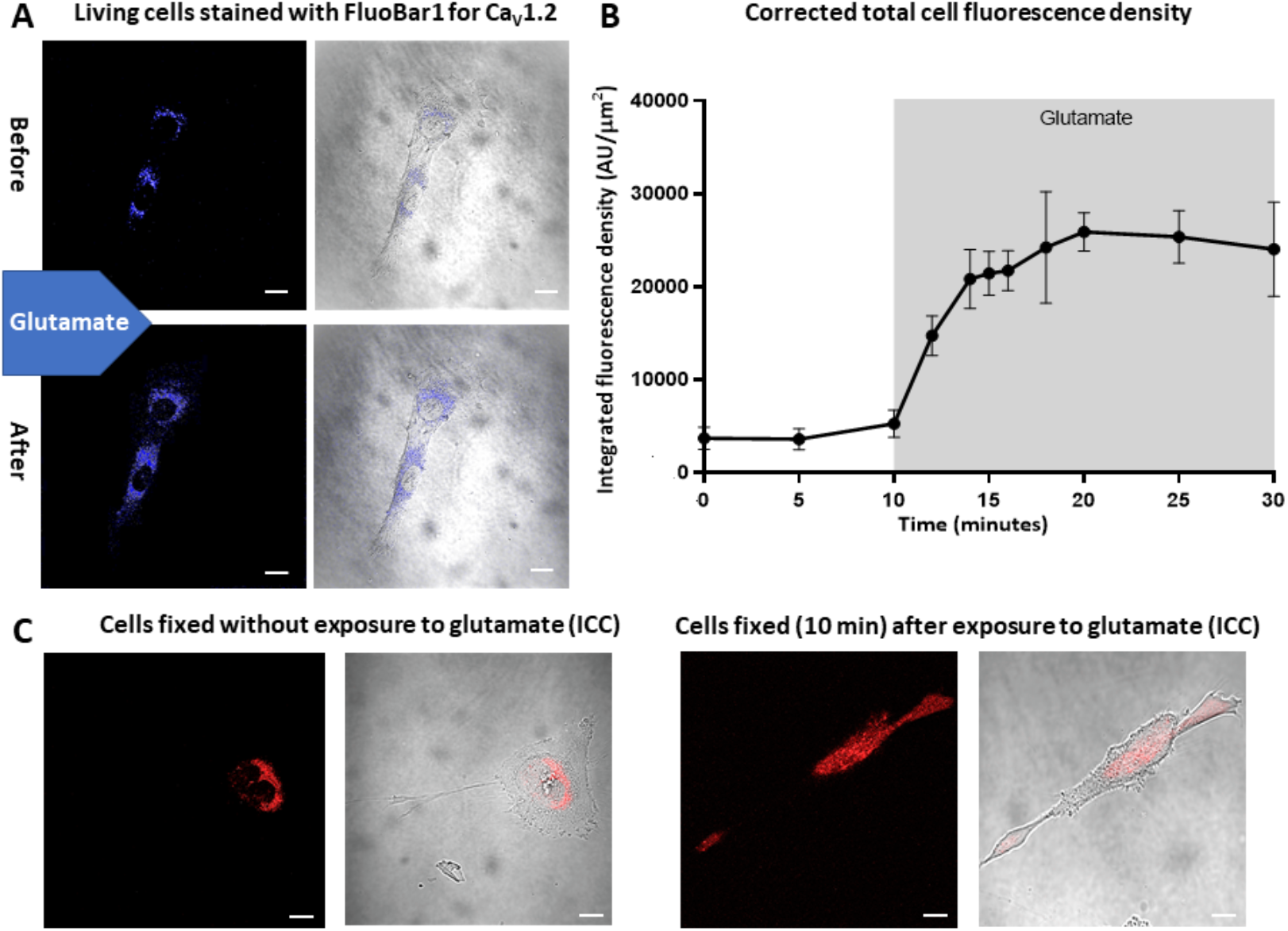
Ca_V_1.2 proteins density and distribution change when astrocyte model cells U118-MG are exposed to 100 μM glutamate. (a) Representative confocal fluorescence micrographs of live cells labelled with 50 nM FluoBar1: prior to glutamate stimulation, and after signal stabilized 20 minutes post-stimulation. (b) Quantified fluorescence from FluorBar1 observed in living cells as a function of time; fluorescence density was measured in each cells and corrected with averaged background density. Two-tailed paired T-test comparison; P value < 0.05, *n* = 3. (c) Representative confocal micrographs of two sets of fixed cells immunostained with an anti-Ca_V_1.2/DyLight550 antibody. U118-MG cells were fixed without stimulation (control), or 10 min after glutamate exposure. All images were acquired at 60X magnification; scale bar is 10 μm.

### Immunocytochemistry labeling shows a change in LTCC distribution patterns following exposure to glutamate

Immunostained U118-MG cells that were *not* exposed to glutamate showed a more localized expression of Ca_V_1.2 channel than cells exposed to 100 μM glutamate for 10 minutes prior to fixation (Fig. 4C). Stimulated cells consistently displayed a widespread distribution and denser exofacial expression of Ca_V_1.2. These observations are static; no before/after information can be obtained from the same cells.

## DISCUSSION

LTCCs are essential ion channel proteins that have four isoforms: CaV1.1, Ca_V_1.2, CaV1.3 and CaV1.4 [49]. In the brain, about 90% of the LTCCs are Ca_V_1.2 and only 10% are CaV1.3 [50]. In neurons, both CaV’s are mostly located at the postsynapse, on the soma, and in the dendritic spines [51]. Despite being "non-excitable" cells, astrocytes also express Ca_V_1.2 and CaV1.3 as their predominant LTCC [47, 52]. The discovery that glutamate can induce a calcium concentration rise in astrocytes sparked research efforts on the complex Ca^2+^-related events in astrocytes that can propagate along cell processes and glia networks [45–48]. To understand astrocytic glutamate signaling, the proteins involved in such dynamic events must be studied in living cells to provide meaningful physiological insights [19].

Recently, we reported that Ca_V_1.2 contributes to the morphological changes induced by glutamate in astrocyte model cells U118-MG [53]. More specifically, Ca_V_1.2 ion channels seem to regulate the extension of filopodia when the astrocytic cell is exposed to glutamate; possibly via its α2δ1 subunit that is responsible for the channel membranar localization [53–58]. A change in Ca_V_1.2 distribution patterns is observable with immunocytochemistry (ICC), but the data must be acquired in *different* sets of cells that are fixed and permeabilized (Fig. 4, C). No simple technique was available to monitor and quantify this alteration in exofacial expression of LTCCs on the *same* cell in real time. At the outset of this study, we hypothesized that a chemical imaging probe would allow us to monitor the real-time redistribution of Ca_V_1.2 in U118-MG cells when they are subjected to a glutamate stimulus.

Imaging probe FluoBar1 (**12,** Fig. 2) was developed to enable the spatiotemporal study of trafficking and/or upregulation of LTCC’s in rapid cellular responses. FluoBar1 is based on unsymmetrical barbiturate **1,** a ligand for LTCCs shown to bind to Ca_V_1.2 and CaV1.3 [36, 37]. Over the past decades, the medicinal chemistry efforts on barbiturates have shown that the apical position of the compounds can be modified without significant impact on the compounds’ activity. Accordingly, we created a 3-steps/one-pot procedure to append a side chain that can be easily attached to any chemical reporter tag for biological studies (e.g., fluorescent tag, biotin, SNAP-tag, etc.). The fluorescent dye Pacific Blue was selected for its small size, high quantum yield, and its distant spectral window from the reference DyLight550-conjugated antibody we used for colocalization studies.

The selectivity of FluoBar1 for LTCC’s is supported by the analysis of its cellular colocalization in cells immunostained with a selective antibody raised against the LTCC isoform Ca_V_1.2 (Fig. 3). Detailed electrophysiology studies on the selectivity of FluoBar1’s parent barbiturate **1** for LTCC’s have been reported by the Striessnig and Soong labs [27, 46]. However, since FluoBar1 was developed for the optical imaging of LTCCs, and not for pharmacological use, this study focused on colocalization analyses as a proof of concept. Labeling the cells five minutes before image acquisition allows one to observe the location of CaV channels in a near-unperturbed state.

FluoBar1 labeled Ca_V_1.2 proteins efficiently; its fluorescence colocalized with immunocytochemical labeling in U118-MG cells that expression Ca_V_1.2 natively (Fig. 3, Table 1) [53, 59]. We also tested whether FluoBar1 can be used as a preliminary screening tool to rapidly detect the presence of LTCCs in a non-denatured cell lysate. This alternative to immuno-dot blotting is much simpler to perform than using antibodies (see ESM for details, Fig S1). In this assay, FluoBar1-dot blotting also showed fluorescence in cells reported to express only CaV1.3 as LTCC (MCF-7) [59]. This observation is consistent with the parent barbiturate **1** being shown to bind to both Ca_V_1.2 and CaV1.3 [36, 37].

Negative controls with COS-7 cells confirmed that (1) no non-selective Ca_V_1.2 labeling occurs with either the FluoBar1 probe or the anti-Ca_V_1.2 antibody; and (2) no passive diffusion of FluoBar1 through the cell membrane occurs within the incubation time (Fig. 3). COS-7 cells are fibroblast-like cells that do not express calcium channels [60, 61]. They were used as heterologous expression system to further validate the probe’s selectivity. The Mender coefficient calculated from colocalization images with CaV.1.2-transfected COS-7 cells suggest that FluoBar1 binds only to properly assembled channels. The epitope for the primary anti-CaV.1.2 antibody used herein binds to a short peptide sequence between second and third internal loop of the channel regardless of the protein’s folding, location, or activity state (residues 848-865, Alomone). In contrast, FluoBar1 can only bind to a properly expressed, folded, and post-translationally modified LTCC [36, 37]. In COS-7 cells, the exogenous expression of the large Ca_V_1.2 voltage-gated protein (α1C) is subject to misfolding and can lead to improper assembly or trafficking [62]. Consequently, it may explain why less fluorescence with FluoBar1 is observed than with anti-Ca_V_1.2 antibody in Ca_V_1.2-transfected COS-7 cells. This is supported by the Mender’s analysis showing 100% colocalization of signals from FluoBar1 (405 nm excitation) with anti-Ca_V_1.2 signals, while only 64% of signals of antibody (559 nm excitation) were colocalized with the probe’s signals (Table 1). A Mender coefficient of 1.00 for the blue channel indicates that all FluoBar1-positive pixels colocalized with an antibody signal. In contrast, the red channel had only a 0.67 coefficient, indicating that the chemical probe colocalized perfectly with the immunological label, but one third of the antibody fluorescence occurred where there was no FluoBar1 signal.

A limitation to this study is that it is not ideal to compare a chemical probe for live cells imaging with immunocytochemistry in fixed cells. A more compelling comparison would be with a recombinant fluorescent protein-fusion to an LTCC; however, CaV’s complex structure do not tolerate well such large changes and would require time-consuming genetic modifications [35]. More work is required to evaluate the selectivity of FluoBar1 for other LTCC’s (e.g., CaV1.1, and CaV1.4).

Finally, real-time imaging of live cells treated with FluoBar1 demonstrated for first time that glutamate causes a prompt redistribution of Ca_V_1.2 in astrocyte model cells U118-MG (Fig. 4A). Glutamate stimulation leads to a rapid 4-fold increase in fluorescence signals density of proteins labelled with FluoBar1 (Fig. 4B). These exciting observations in the *same* cell further support the contribution of LTCC’s in astrocytic glutamate signaling [53]. Access to this imaging tool and simple protocol is now leading us to investigate the dynamics of glutamate-induced process extensions in other glial cells.

## CONCLUSION

In summary, we disclosed FluoBar1 as a new fluorescent probe to visualize LTCCs dynamics in living cells. The synthesis describes how barbiturates can be modified to append a side chain bearing a reporter tag. The labeling selectivity of FluoBar1 for LTCC’s was evaluated by colocalization studies using immunocytochemistry with an anti-Ca_V_1.2 antibody. FluoBar1 was used to monitor the distribution of Ca_V_1.2 in real time, which revealed that Ca_V_1.2 relocalizes within minutes upon glutamate sensing in U118-MG astrocyte cells. This chemical probe provides an alternative to current methods that rely on immunological techniques or molecular biology modifications to monitor protein localization. It also paves the way to investigate the role of LTCC dynamics in calcium events in neurological disorders, e.g., Alzheimer’s, Parkinson’s, and multiple sclerosis [36, 63–67].

## ACKNOWLEDGMENTS

We are grateful to the following UBC researchers for generous gifts: Andis Klegeris for U118-MG cells, Philip Barker for COS-7 cells, and Terry P. Snutch for the pcDNA3-α1c Ca_V_1.2 plasmid.

## AUTHORS CONTRIBUTIONS

MT conceived and performed biological experiments, acquired confocal images, analysed the data, and co-wrote the manuscript. JK conceived and performed the chemical synthesis and characterization. FM conceived the idea and co-wrote the manuscript.

## ETHICAL APPROVAL

This study did not involve protocols requiring ethical approval.

## ELECTRONIC SUPPLEMENTARY MATERIAL

Chemical synthesis procedures, chemical characterization data, additional blotting data.

