## Supplementary Information for "Design of an imaging probe to monitor real-time redistribution of L-type voltage gated calcium channels in astrocytic glutamate signalling"

#### *Electronic Supplementary Information*

### 1. Dot blot assay

Dot blotting (or dot immunobinding) provides a quick way to detect the presence of a protein of interest and/or to assess whether an antibody is not selective (often the secondary antibody).[1] It is a variant of Western blotting in which antigens are detected in samples spotted directly on to a membrane without prior separation.[2] Herein, we compared antibodies and FluoBar1 to selectively label L-type voltage gated calcium channels in a cell lysate—$Ca_V1.2$ and $Ca_V1.3$ in particular.

**Material and Methods**

*Cell culture.* Human astrocytoma U118-MG cells (HTB-15, ATCC), kidney COS-7 cells (CRL-1651™), and breast cancer MCF7 cells (gift from Andrew Jirasek, UBC Okanagan) were cultured in Dulbecco's modified essential medium (DMEM, Gibco 11995-065) supplemented with 10% v/v heat-inactivated fetal bovine serum (HyClone 12483020) 1% v/v penicillin 10,000 U/ml and streptomycin 10,000 µg/ml (Gibco 15140-163). Cells were incubated in a humidified atmosphere containing 5% $CO_2$ at 37 °C until they reached 95% confluency.

*Cell lysate.* Cells were resuspended with prewarmed 0.25% trypsin-EDTA solution (Gibco SH30236.02), then transferred to a 15 mL falcon tube, and spun down at 1000 rpm for 5 min at 4 °C. The pellet was mixed with 50 µL of 1% protease inhibitor cocktail (Amresco, mammalian M250) in sterile Millipore deionized water. Then, 50 µL of NP40 lysis buffer (50 mM Tris-HCl, NaCl 150 mM, 1% NP40-Triton 100x pH 7.4) was added to the solution and incubated for 15 min at 4 °C. The mixture was centrifuged at 3000 rpm for 5 min at 4 °C. The pellet was discarded, and the supernatant was stored at –20 °C.

*Rat brain lysate.* Hippocampal brain tissue from eight weeks rats was homogenized in an ice-cooled buffer containing: Triton100x 1%, 50 mM Tris-HCl (pH 7.4), NaCl 150 mM, EDTA 10 mM, EGTA 10 mM, and 1% protease inhibitor cocktail. The homogenized lysate was centrifuged at 14,000 rpm at 4 °C, the pellet was discarded, and the supernatant was stored at –20 °C until needed.

*Immuno-dot blotting.* Cell lysate aliquots (1 µl) were blotted on two dry nitrocellulose membranes (GE Healthcare, Amersham Protran Sandwich 0.45 µm, GE10600114). The membranes were then blocked for 30 minutes on a rocking plate \using a blocking buffer: 3% bovine serum albumin (BSA, Alfa aesar J64777, New Zealand origin) in 1X Tris-buffered saline + 0.1% Tween (TBS-T). TBS consists Tris 20 mM, NaCl 150 mM. The membranes were washed three times with 1X TBS-T for 5 min each, then followed by incubation with the primary antibody. One membrane was treated with anti-$Ca_V1.2$ primary antibody (rabbit anti- cacna1C, Alomone ACC-003), and the other with anti-$Ca_V1.3$ primary antibody (rabbit anti-cacna1D, Alomone ACC-005); both were incubated for 2 hours at room temperature. The membranes were washed 3 times with TBS-T for 5 minutes each at room temperature on a rocking platform. Both membranes were then incubated with the same secondary antibody (goat anti-rabbit IgG HRP-conjugated, HAF008) for one hour at room temperature. Membranes were washed three times with TBS-T each for 5 minutes; the blots were visualized by a 1:1 solution of luminol + enhancer (Pierce ECL Western blotting substrate, 32106) using a quantitative Western blot imaging system (Alpha-Innotech FluorChem HD2).

[1] Tian G, Tang F, Yang C, Zhang W, Bergquist J, Wang B, Mi J, Zhang J, *Oncotarget.* **2017.** 8:58553–58562 . https://doi.org/10.18632/oncotarget.17236
[2] Stott DI *J. Immunol. Methods.* **1989.** 119:153–187

*Fluobar1 dot-labeling.* Cell lysate aliquots (1 µl) were blotted on a dry nitrocellulose membrane. The dried membrane was soaked in a 50 nM FluoBar1 solution in sterile millipore dionized water for 1 minute. The membrane was rinsed twice with sterile millipore deionized water, then immediately placed on a fluorescence confocal microscope slide and imaged at 10x magnification and under laser excitation at 405 nm (Olympus FV1000, AlexaFluor-405 filter). The membrane was kept wet during the imaging processes.

**Results**

Immunoblotted membranes show the presence of $Ca_V1.2$ in U118-MG cells and brain lysate, but not in MCF7 cells. (Fig. S1-a). Presence of $Ca_V1.3$ is observed in MCF7 cells and in rat brain lysate, and only at trace levels in U118-MG cells. COS-7 cells are a negative control as they do not express L-type $Ca_V$'s to a detectable level. Membranes stained with the imaging probe FluoBar1 show fluorescence in both MCF7 cells and U118-MG cells when irradiated at 405 nm, suggesting that FluoBar1 binds to $Ca_V1.2$ and $Ca_V1.3$ proteins (Fig. S1-b). Brain lysates also show robust fluorescence. Only background levels of fluorescence were present with COS-7 cells, suggesting that minimal unselective binding occurs with FluoBar1 (Figure S1-b).

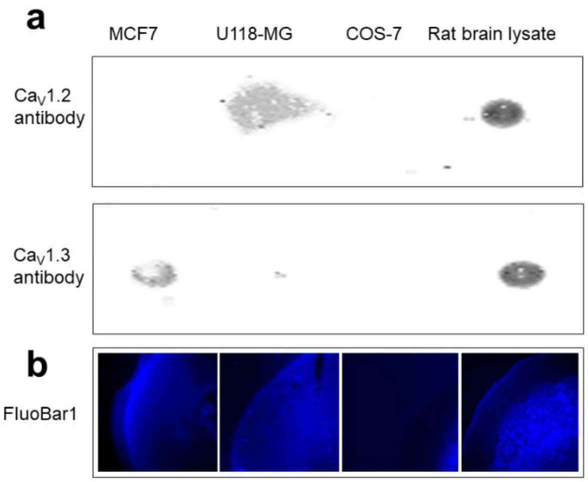


**Figure S1.** FluoBar1 shows affinity for both $Ca_V1.2$ and $Ca_V1.3$. (a) Immuno-dot blotting of cell lysates from MCF7 cells, U118-MG cells, COS-7 cells and brain lysate by chemiluminescence. (b) FluoBar1 dot-labeling imaged by fluorescence confocal microscopy, 10x magnification.

Immuno-dot blotting is a crude alternative to Western blotting that bypasses electrophoresis separation, yet relies on antibodies and requires a workflow typically spanning a few hours. In this study, we used our new chemical probe, FluoBar1, to detect the presence of L-type voltage gated calcium channel on a blotted membrane using a 1 minute incubation time.

We selected cell lines with different $Ca_V$ protein expression profiles to evaluate the preliminary selectivity of FluoBar1 for LTCCs. U118-MG cells naturally express $Ca_V1.2$, MCF7 cells express $Ca_V1.3$,[3] and COS-7 cells are calcium-channel free cells.[4,5] FluoBar1 did not show significant non-selective labeling with LTCC-free cell lysates. The parent molecule of FluoBar1, barbituate **1** (Fig. S2), was shown to bind both to $Ca_V1.2$ and $Ca_V1.3$.[5,6]

[3] Thul PJ, et al., *Science* . **2017.** 356:eaal3321 . https://doi.org/10.1126/science.aal3321
[4] Xu W, Lipscombe D. *J Neurosci* . **2001.** 21:5944–5951 . https://doi.org/21/16/5944 [pii]
[5] Lao QZ, Kobrinsky E, Liu Z, Soldatov NM, *FASEB J* . **2010**. 24:5013–5023 . https://doi.org/10.1096/fj.10.165381
[6] Ortner NJ, Bock G, Vandael DHF, Mauersberger R, Draheim HJ, Gust R, Carbone E, Tuluc P, Striessnig. *J. Nat Commun.* **2014**. 5: https://doi.org/10.1038/ncomms4897

### 2. Chemical Synthesis

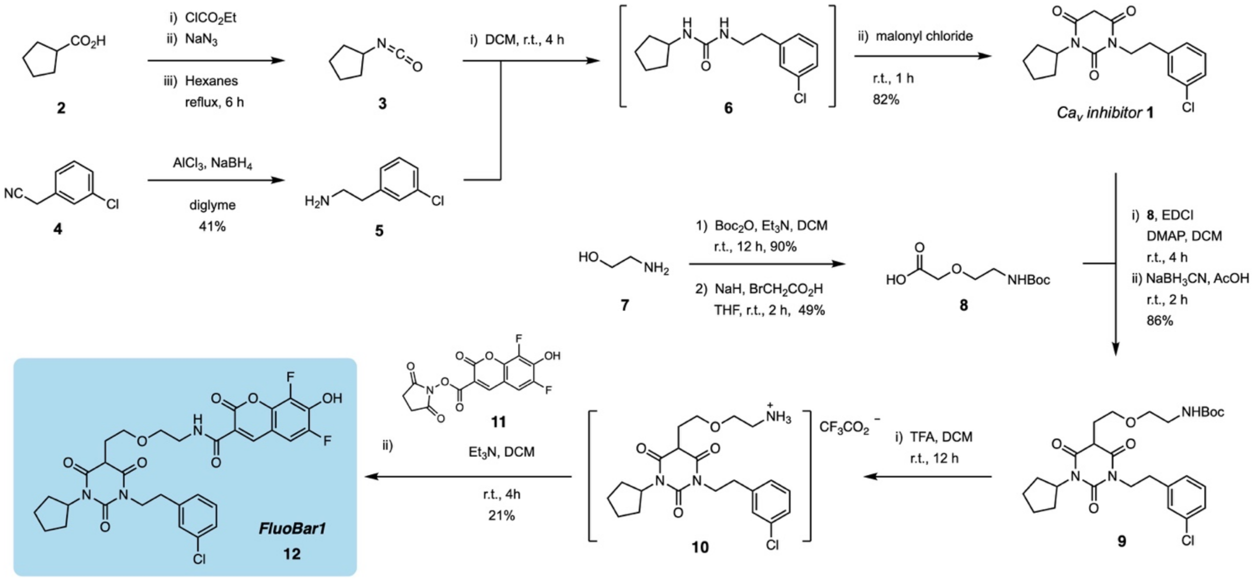


**Figure S2.** Complete synthetic route to obtain imaging probe FluoBar1.

**General Synthesis Procedures.** All non-aqueous reactions were carried out under a nitrogen or argon atmosphere, in flame-dried single-neck, round bottom flasks fitted with a rubber septum and with magnetic stirring. Air or water sensitive liquids and solutions were transferred via syringe or stainless-steel cannula. Organic solutions were concentrated by rotary evaporation at 25–45 °C at 50-200 torr. Thin layer chromatography (TLC) was performed on glass plates precoated with Silica gel F254, 250 $\mu$m, 60 Å, from EMD Chemicals Inc (EMD 5715-1). TLC plates were visualized under a 254 or 365 nm UV light source, then stained by chemical reagent: typically, iodine vapors were used first, followed by one of the following solutions: acidic ethanolic vanillin or basic aq. potassium permanganate. The TLC plate was then heated briefly to 200 ºC using a heat gun. Column chromatography purifications were performed with 230-400 mesh silica gel from Silicycle, Quebec (SiliaFlash R12030B, P60, 40-63 $\mu$m, 60 Å).

**Materials.** Reagents and starting materials were purchased from: Sigma-Aldrich, Oakwood Chemicals, Alfa Aesar, Acros Organics, TCI America, or Fisher Scientific and were used as received unless otherwise noted. Succinimidyl 6,8-difluoro-7-hydroxycoumarin-3-carboxylate (**11**) was synthesized as described previously.[7] Tetrahydrofuran, dichloromethane, hexanes, toluene, and diethyl ether were purified on a glass contour solvent purification system under an argon atmosphere.[8] $Et_3N$ and pyridine were distilled from $CaH_2$. Solvents used for chromatographic purifications were obtained from Fischer Scientific or VWR and used without further purification.

**Instruments.** $^1$H, $^{13}$C, and $^{19}$F NMR spectra were recorded on a 400 MHz Varian NMR AS400 equipped with an ATB-400 probe at 25 °C. NMR spectra were analyzed with MestReNova 10 from Mestrelab Research. Proton chemical shifts are expressed in ppm ($\delta$ scale) downfield from tetramethylsilane and are referenced to this standard. Carbon chemical shifts are expressed in ppm ($\delta$ scale) downfield from tetramethylsilane and are referenced to carbon resonance of the NMR solvent ($CDCl_3$ $\delta$ 77.00, *d*$_6$-DMSO $\delta$ 39.52) or MeOH $\delta$ 49.50 in case of $D_2O$. Fluorine chemical shifts are expressed in ppm ($\delta$ scale) and referenced to $FCCl_3$ (0.0 ppm). Spectral data are listed as follows: chemical shift,

[7] Kerkovius, J.K., and Menard, F. (2016). A practical synthesis of 6,8-difluoro-7-hydroxycoumarin derivatives for fluorescence applications. *Synthesis 48*, 1622.

[8] Pangborn, A.B., Giardello, M.A., Grubbs, R.H., Rosen, R.K., and Timmers, F.J. (1996). Safe and convenient procedure for solvent purification. *Organometallics 15*, 1518.

integration, multiplicity (s = singlet, d = doublet, t = triplet, q = quartet, m = multiplet, br s = broad singlet), and coupling constant (*J*, Hz). Infrared spectra (IR) of thin films were obtained using a Spectrum Two FT-IR spectrometer (Perkin-Elmer), and the characteristic absorptions are given in wavenumbers. High-resolution mass spectra were obtained in ESI mode using an HCTultra PTM discovery system spectrometer (Mass Spectrometry Facility, UBC Vancouver). Melting points of solid samples were measured with an IA9200 melting point apparatus (Electrothermal) or micro-melting point apparatus Mel-TempII (Laboratory Devices, USA). The final product was purified by HPLC Waters Delta Prep system on a C18 column (Nova-Pak® HR C18, 19×300 mm, 6 $\mu$m, Waters) with fluorimetric detector.

---

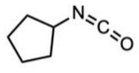

**Cyclopentyl isocyanate (3).**[9] To a 100 mL flask was added cyclopentane carboxylic acid (12.2 g, 107 mmol), hexane (75 mL) and triethylamine (15.4 mL, 110 mmol). To a 500 mL flask was added hexane (100 mL) and ethyl chloroformate (10.2 mL, 107 mmol). The ethyl chloroformate solution was cooled on an ice bath to 0 °C. The cyclopentane carboxylic acid solution was added dropwise via a cannula to the ethyl chloroformate solution over the course of 30 minutes. Once the addition was complete, the white slurry was stirred for 30 minutes on ice, vacuum filtered, and washed once with hexanes (30 mL). The filtrate was transfered to a 500 mL flask and was concentrated under reduced pressure to a volume of about 20 mL. To the concentrated solution was added acetone (100 mL), then equiped with an addition funnel containing a solution of sodium azide (11.0 g, 169 mmol) in water (100 mL). The acetone solution was cooled on an ice bath and the sodium azide solution was added dropwise over the course of 15 minutes. The reaction was stirred for an additional 2 hours on ice. The reaction was poured into ice cold water (250 mL) and extracted with hexane (2 x 125 mL). The organic layer was dried over anhydrous $MgSO_4$, filtered and added to a 500 mL flask. The hexane solution was refluxed under nitrogen for 4 hours, and then concentrated under reduced pressure to a volume of 20-40 mL. The crude product was estimated to be a 43% solution in hexanes as determined by $^1$H NMR in $CDCl_3$ (14.75 g of solution, est. 6.34 g of product). The crude product in solution was used as is for the next step.

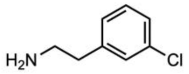

**2-(3-Chlorophenyl)ethylamine (5).**[10] A 500 mL flask was charged with diglyme (260 mL) and sodium borohydride (9.50 g, 251 mmol). To the sodium borohydride suspension was added 2-(3-chlorophenyl)acetonitrile (10.0 g, 66.0 mmol). To a 250 mL Erlenmeyer flask was added diglyme (60 mL) followed by a slow addition of anhydrous aluminum chloride (11.15 g, 83.6 mmol) at –78 ºC. *Warning:* Addition of $AlCl_3$ to diglyme is extremely exothermic. The aluminum trichloride suspension was added to the sodium borohydride and nitrile solution over the course of 20 minutes. Gas evolution occurred, and the reaction turned lemon yellow. The reaction was stirred at r.t. for 4 hours. The reaction was quenched by a slow addition to a mixture of 12 M HCl (86 mL) in ice (200 g) with simultaneous external cooling on an ice bath. The quenched solution was extracted with DCM (3 x 150 mL). The combined organic layer was washed with water (2 x 100 mL), dried over anhydrous $MgSO_4$, and filtered. The solution was concentrated under reduced pressure to yield a brown oil. To the crude product (which is obtained as a solution in diglyme). To the product was added oxalic acid dihydrate (5.0 g, 1.05 eq) as a solution in water (100 mL). The solution was heated to boiling, and was gravity filtered to remove an insoluble green residue. The solution was cooled to r.t. and then to 0 ºC. The white crystalline cake was broken up, vacuum filtered, and washed with ice cold water (3 x 30 mL). The solids were added to water (200 mL) in a seperatory funnel. To the aqueous suspension was added 40% NaOH added until the pH was above 11. The mixture was extracted with diethyl ether (2 x 150 mL). The combined organic layer was copiously washed with water (8 x 75 mL), brine (75 mL), dried over anhydrous $MgSO_4$, and filtered. The solvent was removed under reduced pressure to yield a clear and nearly colourless oil. (4.22 g, 41%). Characterization data matched literature.[11] **$^1$H NMR** (400 MHz, $CDCl_3$) $\delta$ = 7.19 (m, 4H), 2.95 (t, $J$ = 7.03 Hz, 2H), 2,71 (t, $J$ = 7.03 Hz, 2H), 1.35 (br s, 2H) ppm. **$^{13}$C NMR** (101 MHz, $CDCl_3$) $\delta$ = 142.1, 134.3, 129.8, 129.0, 127.2, 126.5, 43.4, 39.8 ppm. **IR** (FT-IR, neat film): 3380, 3285, 3059, 2929 cm$^{-1}$.

---

[9] Eda, M., Takemoto, T., Ono, S.I., Okada, T., Kosaka, K., Ghoda, M., Matzno, S., Nakamura, N., Fukaya, C. *J. Med. Chem.* **1994**. *37*, 1983-1990.
[10] Brown, H.C., Rao, S.B.C. *J. Am. Chem. Soc.* **1955**. *78*, 2582-2588.
[11] https://www.sigmaaldrich.com/spectra/fnmr/FNMR007463.PDF

**1-(3-Chlorophenethyl)-3-cyclopentylpyrimidine-2,4,6-trione (1).**[12] To a 500 mL flask was added 2-(3-chlorophenyl)ethylamine (**5**) (1.116 g, 7.17 mmol) and dichloromethane (220 mL). To the dichloromethane solution was added cyclopentyl isocyanate **3** as a 7% solution in hexanes (10.4 g of solution, 0. 728 g isocyanate, 6.55 mmol) over the course of 5 minutes. The reaction was stirred at r.t. until complete by TLC (50% EtOAc in hexanes, about 1 hour). Malonyl chloride (0.77 mL, 7.92 mmol) was added dropwise to the reaction solution over the course of 5 minutes. The reaction was stirred for 1 hour at r.t. after the malonyl chloride addition was complete. The reaction was washed with brine (50 mL), dried over anhydrous $MgSO_4$, filtered and concentrated under reduced pressure to yield a yellow solid. The crude product was purified by flash chromatography on silica gel (20% EtOAc in hexanes). The product was obtained as a white powder (1.97 g, 82%). Characterization data matched literature.[12] **$^1$H NMR** (400 MHz, $CDCl_3$) $\delta$ = 7.22 (m, 4H), 5.14 (m, 1H), 4.08 (m, 2H), 3.62 (s, 2H), 2.88 (m, 2H), 1.84-1.94 (m, 6H), 1.57 (m, 2H) ppm. **$^{13}$C NMR** (101 MHz, $CDCl_3$) $\delta$ = 164.9, 164.7, 151.1, 140.0, 134.5, 130.0, 129.3, 127.3, 127.2, 54.6, 42.8, 40.3, 33.9, 28.9, 25.7 ppm. **IR** (FT-IR, solid state): 3063, 2959, 1673 cm$^{-1}$.

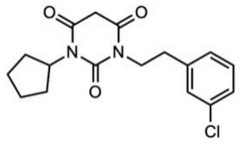


***tert*-Butyl-(2-hydroxyethyl)carbamate (7b).**[13] To a 250 mL flask was added ethanolamine (2.51 g, 41.0 mmol), DCM (125 mL), and triethylamine (11.4 mL, 81.2 mmol). The solution was cooled on an ice bath to 0 ºC. A solution of $Boc_2O$ (10.08 g, 46.2 mmol) in DCM (15 mL) was added dropwise to the amine solution over the course of 10 minutes. The reaction was stirred overnight at r.t. The reaction was washed with saturated ammonium chloride (3 x 125 mL), dried over anhydrous $MgSO_4$, filtered and concentrated under reduced pressure to yield the crude product as a yellow oil. The oil was purified by flash chromatography on silica gel (50% EtOAc in hexanes). The product was isolated as a clear colourless oil (11.66 g, 88%). Characterization data matched literature precedents.[13,14] **$^1$H NMR** (400 MHz, $CDCl_3$) $\delta$ = 5.31 (br s, 1H), 3.67 (m, 2H), 3.59 (br s, 1H), 3.26 (m, 2H), 1.44 (s, 9H) ppm. **$^{13}$C NMR** (101 MHz, $CDCl_3$) $\delta$ = 157.2, 80.0, 62.7, 43.5, 28.8 ppm. **IR** (FT-IR, neat film): 3337, 2981, 1694 cm$^{-1}$.

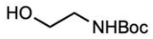


**2-(2-((*tert*-Butoxycarbonyl)amino)ethoxy)acetic acid (8).**[15] To a 250 mL flask was added *tert*-butyl-(2-hydroxyethyl)carbamate (4.64 g, 28.8 mmol), bromoacetic acid (2.02 g, 14.5 mmol) and THF (50 mL). The solution was cooled to 0 ºC on an ice bath. 60% NaH in mineral oil (1.73 g, 43.4 mmol) was added in portions to the reaction over the course of 5 minutes. The reaction was stirred at r.t. for 2 hours after the NaH addition was complete. The reaction was poured into water (300 mL) and extracted with diethyl ether (3 x 200 mL). The aqueous layer was acidified with 1 M HCl until it was below pH 3. The acidified aqueous layer was extracted with diethyl ether (3 x 200 mL). The organic extracts of only the acidified aqueous layer were combined and were washed with saturated sodium bicarbonate (2 x 100 mL), brine (100 mL), dried over anhydrous $MgSO_4$, filtered and concentrated under reduced pressure. The crude product was obtained as a pale yellow oil which was purified by flash chromatography on silica gel (washed with 10% acetic acid in EtOAc) eluting with 50% EtOAc in hexanes. The product was obtained as a clear colourless oil (2.66 g, 83%). Characterization data matched commercial source.[16] **$^1$H NMR** (400 MHz, $CDCl_3$) $\delta$ = 10.7 (br s, 1H), 5.22 (br s, 1H), 4.09 (s, 2H), 3.58 (t, $J$ = 5.1 Hz, 2H), 3.30 (br s, 2H), 1.40 (s, 9H) ppm. **$^{13}$C NMR** (101 MHz, $CDCl_3$) $\delta$ = 174.4, 156.6, 79.9, 71.0, 68.1, 40.5, 28.5 ppm. **IR** (FT-IR, neat film): 3350-2985, 1708 cm$^{-1}$.

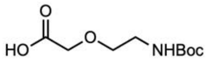


[12] Kang, S., Cooper, G., Dunne, S.F., Luan, C.H., Surmeier, J.D., Silverman, R.B. *J. Med. Chem.* **2013**. *56*, 4786-4797.
[13] Mansueto, M., Frey, W., Laschat, S. *Chem. Eur. J.* **2013**. *19*, 16058-16065.
[14] Snead, A.N., Miyakawa, M., Tan, E.S., Scanlan, T.S. *Bioorg. Med. Chem. Lett.* **2008**, *18*, 5920-5922.
[15] Knoller, H., Heckmann, D., Hacket, F., Zander, N., Nocken, F. WO 2012/004006 Al, January 12, 2012.
[16] Miilipore Sigma 901579. https://www.sigmaaldrich.com/catalog/product/aldrich/901579

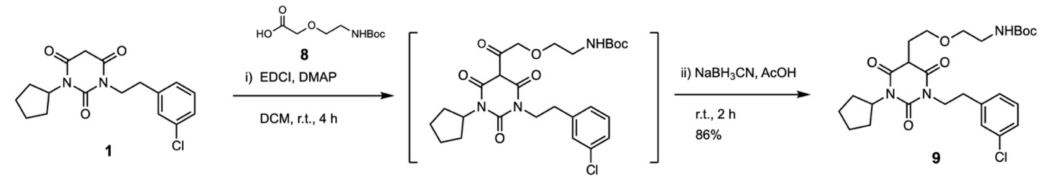

***tert*-Butyl-(2-(2-(1-cyclopentyl-2,4,6-trioxo-3-phenethyl-hexahydropyrimidin-5-yl)ethoxy)ethyl)carbamate (9).**

This procedure was adapted from the literature.[17] To a 100 flask was added the barbiturate **1** (0.501 g, 1.50 mmol), 2-(2'-(*N*-Boc-amino)ethoxy)acetic acid (**8**) (0.406 g, 1.85 mmol), DMAP (0.101 g, 0.827 mmol) and DCM (30 mL). The solution was cooled to 0 °C on ice. To the solution was added EDC (0.317 g, 1.65 mmol) in a single portion as a solid. The reaction was stirred on ice for 3 minutes and then at r.t. for 3 hours. The reaction mixture was concentrated under reduced pressure to yield a thick yellow oil. The oil was dissolved in 8 mL of glacial acetic acid. To the acetic acid solution was added solid sodium cyanoborohydride (0.291 g, 4.63 mmol) in a single portion in air. The reaction was stirred for a 2 hours at r.t. once the sodium cyanoborohydride had been added. The reaction mixture was diluted in water (20 mL) and extracted with DCM (3 x 20 mL). The combined organic extracts were dried over anhydrous MgSO4, filtered and concentrated under reduced pressure. The crude oil was purified by flash chromatography on silica gel (5% to 20% $Et_2O$ in DCM gradient). The residual solvents were removed from the product in vacuo (0.1 torr) for 24 hours. A clear colourless viscous oil was obtained (0.670 g, 86%). **$^1$H NMR** (400 MHz, $CDCl_3$) δ = 7.22 (m, 4H), 5.15 (m, 1H), 4.79 (br s, 1H), 4.08 (m, 2H), 3.49 (m, 3H), 3.38 (t, J = 5.3 Hz, 2H), 3.18 (q, J = 4.7 Hz, 2H), 2.87 (t, J = 7.8 Hz, 2H), 2.44 (m, 2H), 1.81-1.99 (m, 6H), 1.54-1.63 (m, 2H), 1.41 (s, 9H) ppm. **$^{13}$C NMR** (101 MHz, $CDCl_3$) δ = 175.2, 168.6, 168.4, 156.1, 151.0, 140.2, 134.5, 130.0, 129.3, 127.4, 127.1, 79.6, 70.4, 67.1, 54.6, 53.6, 46.4, 42.8, 40.4, 33.9, 30.1, 29.0, 28.8, 28.5, 25.8, 20.8 ppm. **IR** (FT-IR, thin film): 3380, 2955, 1703 $cm^{-1}$. **HRMS** (ESI-TOF) *m/z*: $[M+Na]^+$ calcd for $C_{26}H_{36}ClN_3NaO_6^+$ 544.2185; found 544.2191.

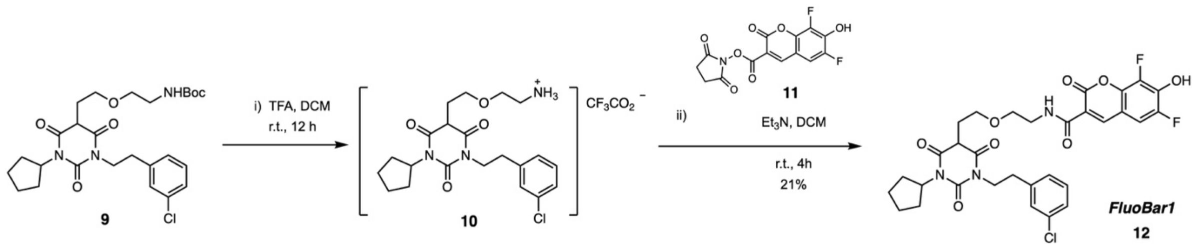

***N*-(2-(2-(1-(3-Chlorophenethyl)-3-cyclopentyl-2,4,6-triox-ohexahydropyrimidin-5-yl)ethoxy)ethyl)-6,8-difluoro-7-hydroxycoumarin-3-carboxamide — *FluoBar1* (12).**

To a 250 mL flask was added the linked barbiturate **9** (0.386 g, 0.741 mmol), and DCM (40 mL). Trifluoroacetic acid (1.2 mL, 15.67 mmol) was added to the solution at r.t. and the reaction was stirred overnight. The next day, the flask was protected from light, triethylamine (4.2 mL, 30.1 mmol) was added followed immediately by a solution of the NHS ester of Pacific Blue **11** (0.261 g, 0.765 mmol) in DMF (15 mL). The reaction was stirred for 4 hours at r.t. The reaction was quenched by slow addition to water (50 mL), followed by 1 M HCl aq. solution until the solution was below pH 4. The mixture was extracted with ethyl acetate (3 x 40 mL). The combined organic layers were washed with water (2 x 40 mL), brine (50 mL), dried over anhydrous $MgSO_4$, filtered and concentrated under reduced pressure. The crude brown oil was purified by flash chromatography on silica gel (70% EtOAc in hexanes). The semi-purified product was a yellow powder and was further purified by HPLC. A 50 mg/mL solution of the product was prepared in acetonitrile. Purification was performed on an Agilent 1290 infinity UPLC with a Waters Nova-Pak® HR C18 6 µm 19x300 mm prep HPLC column. The sample was loaded onto the column in 160 µL aliquots and eluted with an isocratic mixture of 70% acetonitrile containing 0.1% formic acid and 30% water containing 0.2% formic acid at a rate of 3.000 mL/min. The

[17] Szostak, M., Sautier, B., Spain, M., Behlendorf, M., Procter, D.JT. *Angew. Chem. Int. Ed.* **2013**. *52*, 12559-12563.

product was detected at 365 nm on a diode array detector. The product eluted at 35 minutes. The collected fractions were lyophilized to yield a light yellow/green powder (93 mg, 21%).

The purity of each fraction was analyzed using 2 μL injection onto a Waters Nova-Pak® C18 4 μm bead 3.9x300 mm analytical column eluting with an isocratic mixture of 80% acetonitrile containing 0.1% formic acid and 20% water containing 0.2% formic acid at 0.3 mL/min. The sample was detected at 365 nm; purity was determined by comparing relative integration of the product to impurity peaks. An NMR sample was prepared by dissolving the crude in an excess of CDCl3 followed by filtration through a 20–40 μm syringe filter to remove traces of an insoluble residue. The solution was concentrated to a volume of 0.7-1.0 mL and loaded into an NMR tube. The product is sensitive: it was found to degrade over time with noticeable impurities appearing after 24 hours at r.t. **$^{1}$H NMR** (400 MHz, $CDCl_3$) δ = 8.84 (br s, 1H), 8.69 (d, $J$ = 1.4 Hz, 1H), 7.21-7.11 (m, 5H), 5.13 (p, $J$ = 8.5 Hz, 1H), 4.06 (td, $J$ = 7.8, 4.1 Hz, 2H), 3.59 (m, 7H), 2.85 (t, $J$ = 7.4 Hz, 2H), 2.40 (m, 2H), 2.00-1.79 (m, 6H), 1.61-1.51 (m, 2H) ppm. **$^{13}$C NMR** (101 MHz, $CDCl_3$) δ =

168.7, 168.5, 161.7, 160.1, 151.0, 148.7 (dd, $J$ = 244.0, 3.6 Hz), 147.6, 140.8 (dd, $J$ = 9.2, 1.7 Hz), 140.1, 139.6 (dd, $J$ = 18.4, 12.8 Hz), 138.8 (dd, $J$ = 249.7, 5.8 Hz), 134.4, 129.9, 129.2, 127.2, 126.7, 116.7, 110.5 (d, $J$ = 9.4 Hz), 110.0 (dd, $J$ = 20.2, 3.5 Hz), 69.2, 67.3, 54.7, 46.2, 42.9, 39.8, 33.8, 30.1, 28.9, 28.8, 25.8, 25.7 ppm. **$^{19}$F NMR** (377 MHz, CDCl3) δ = –136.2 (t, $J$ = 9.2 Hz), –152.7 (d, $J$ = 8.8 Hz) ppm. **IR** (FT-IR, solid state): 3345, 3060, 2948, 2866, 1720, 1676 cm$^{-1}$. **HRMS** (ESI-TOF) *m/z*: [M–H]$^-$ calcd for $C_{31}H_{29}ClF_2N_3O_8$ 644.1616; found 644.1619.

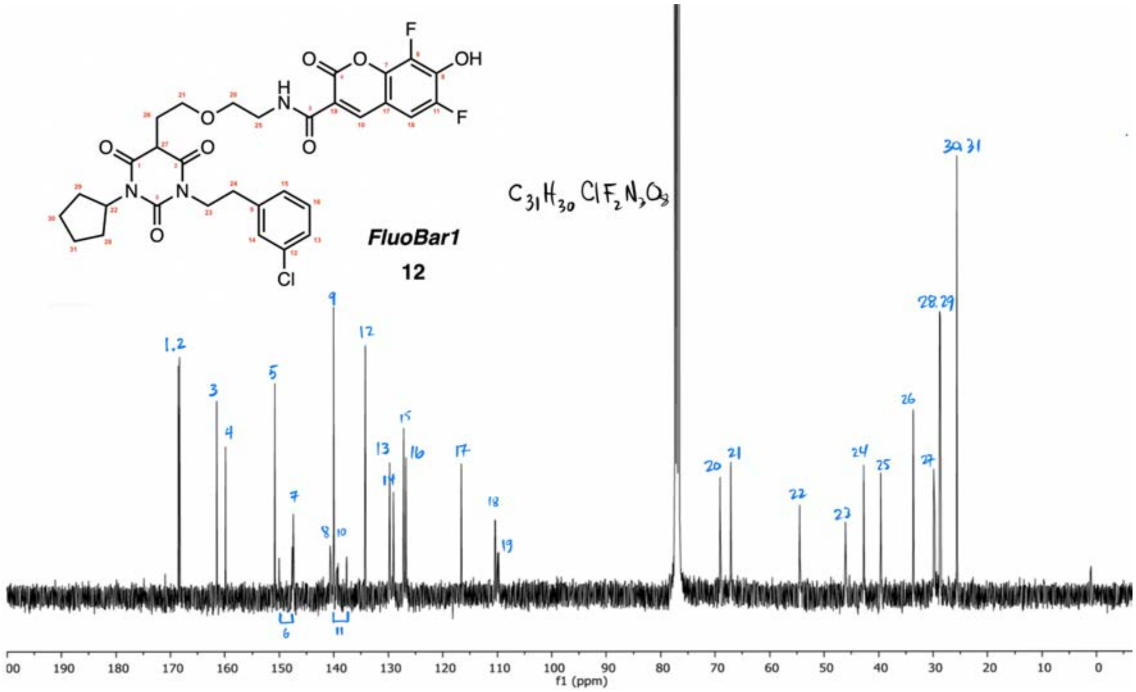

### 3. Characterization Spectra

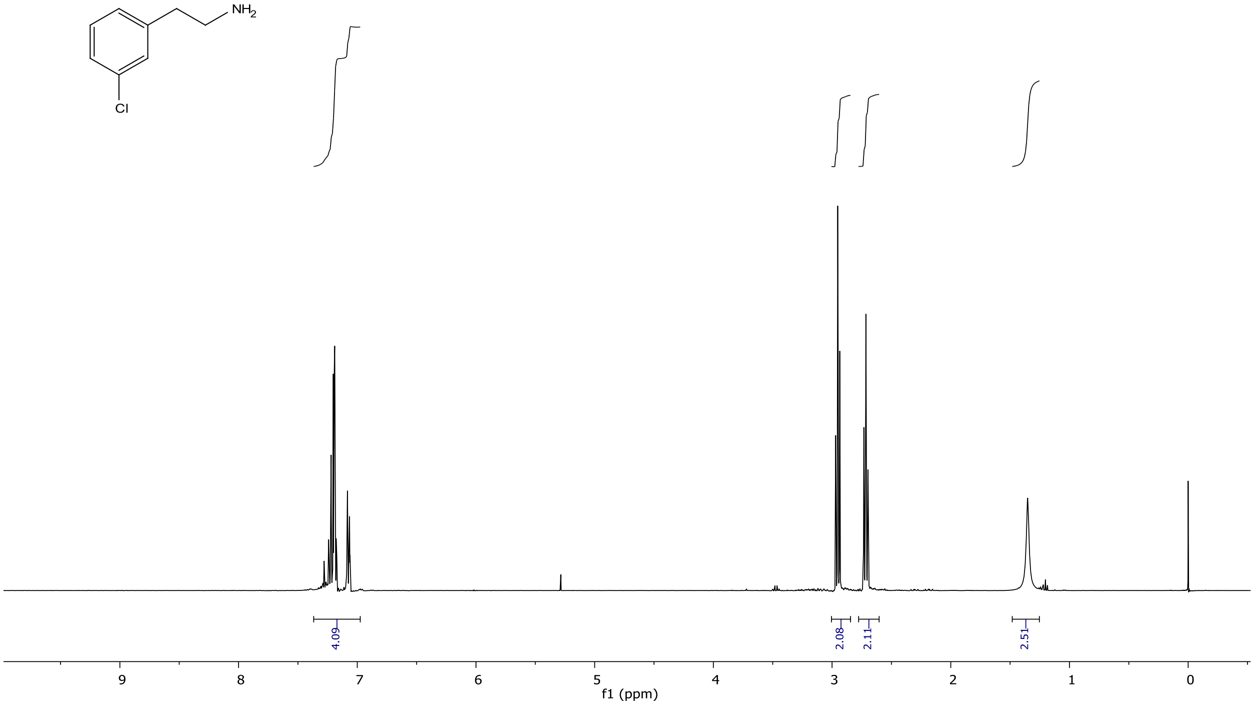


**Figure S3.** **¹H NMR** (400 MHZ, $CDCl_3$): 2-(3-C7hlorophenyl)ethylamine (**5**)

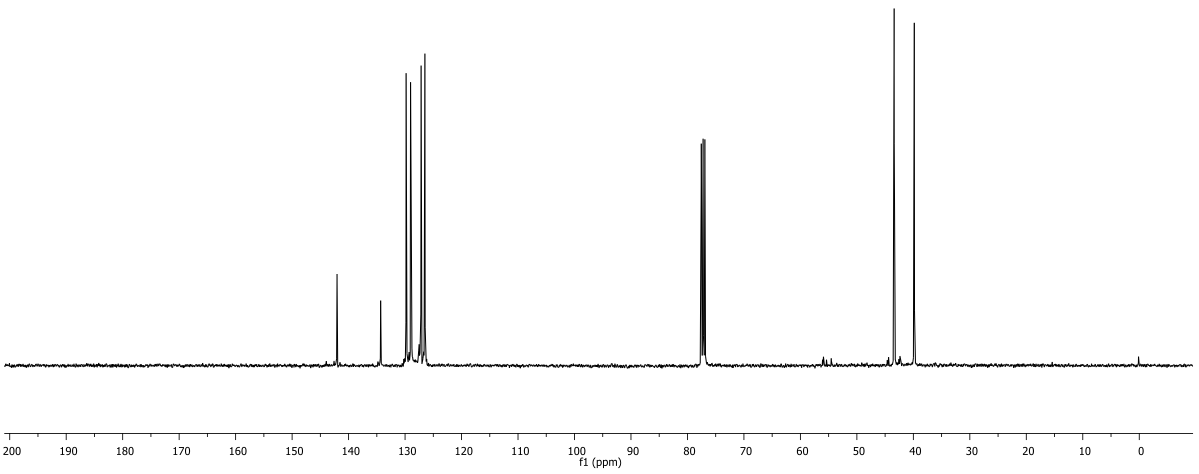


**Figure S4.** **¹³C NMR** (101 MHZ, $CDCl_3$): 2-(3-Chlorophenyl)ethylamine (**5**)

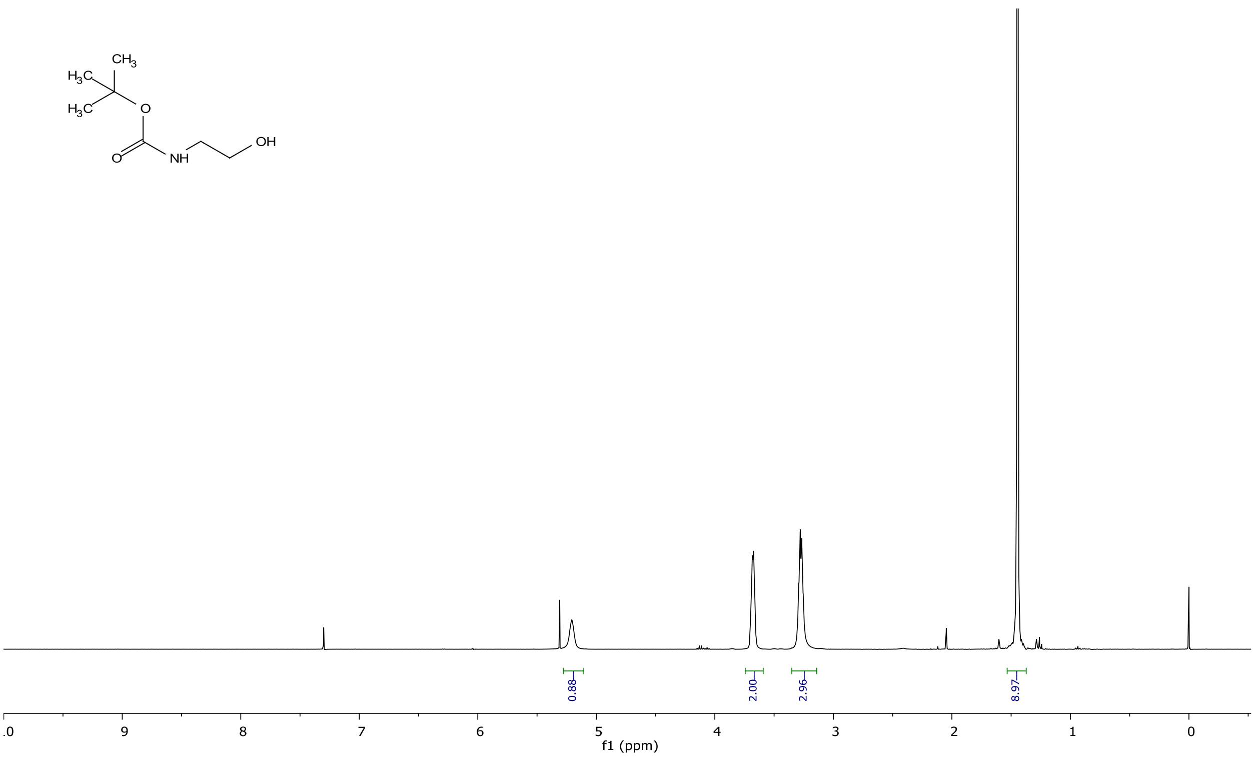


**Figure S5.** **$^{1}$H NMR** (400 MHz, $CDCl_3$): *tert*-Butyl-(2-hydroxyethyl)carbamate (**7b**)

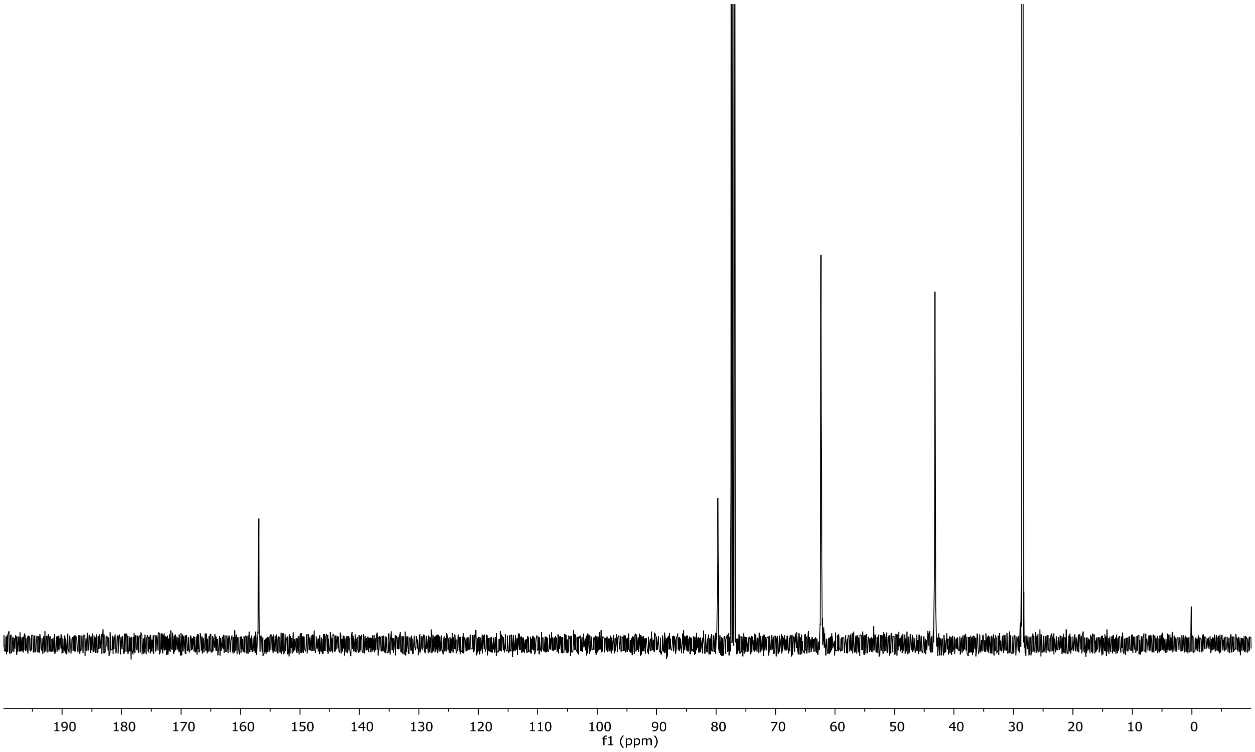


**Figure S6.** **$^{13}$C NMR** (101 MHZ, $CDCl_3$): *tert*-Butyl-(2-hydroxyethyl)carbamate (**7b**)

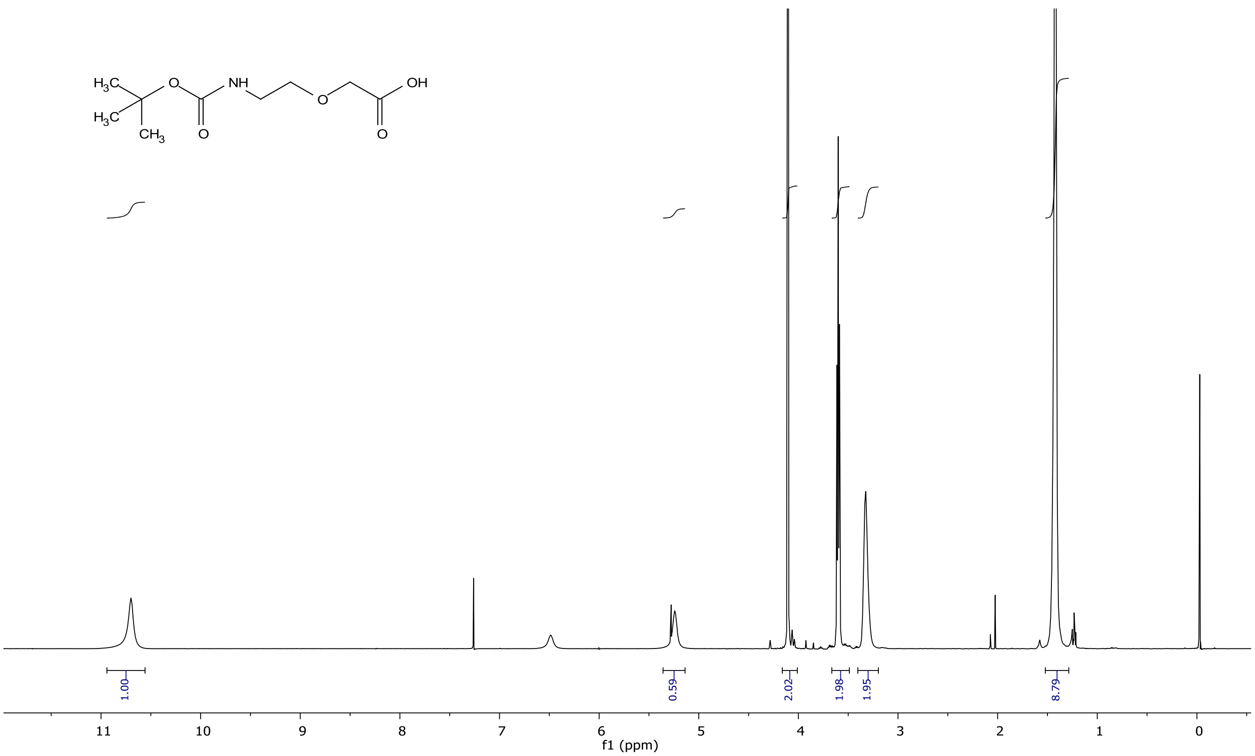


**Figure S7**. **$^{1}$H NMR** (400 MHz, $CDCl_3$): 2-(2-((*tert*-Butoxycarbonyl)amino)ethoxy)acetic acid **(8)**

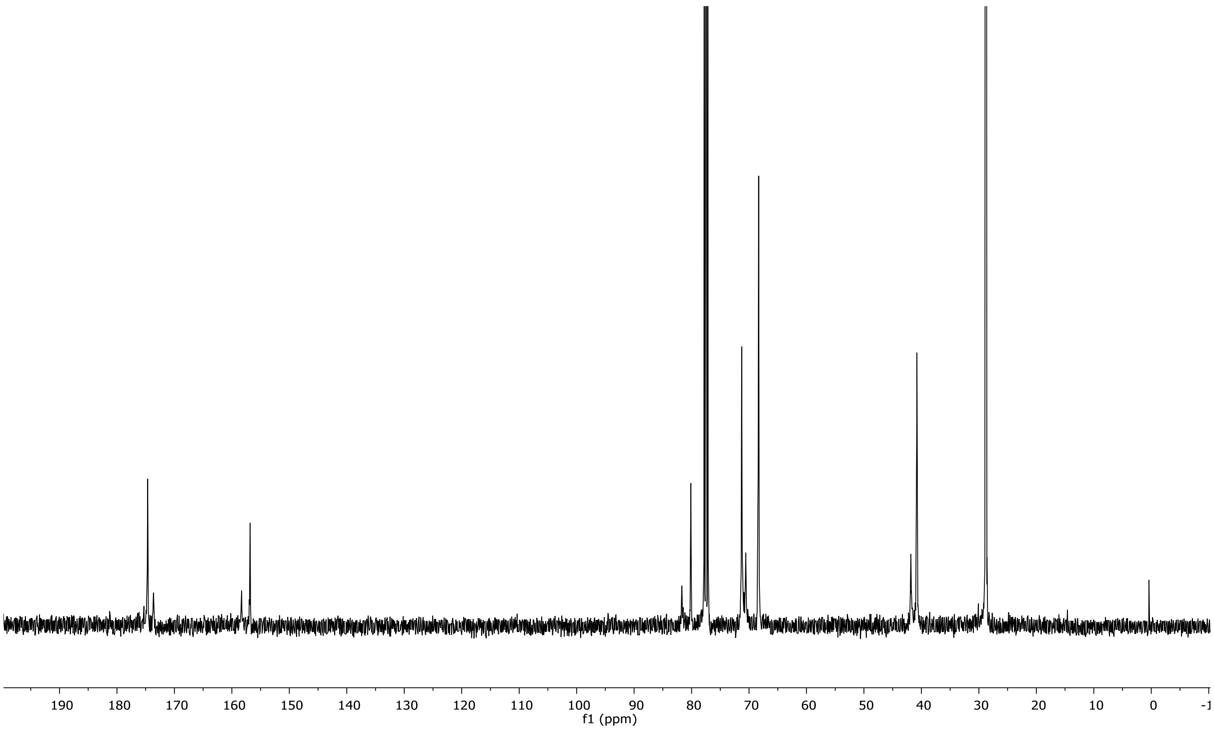


**Figure S8.** **$^{13}$C NMR** (101 MHZ, $CDCl_3$): 2-(2-((*tert*-Butoxycarbonyl)amino)ethoxy)acetic acid **(8)**

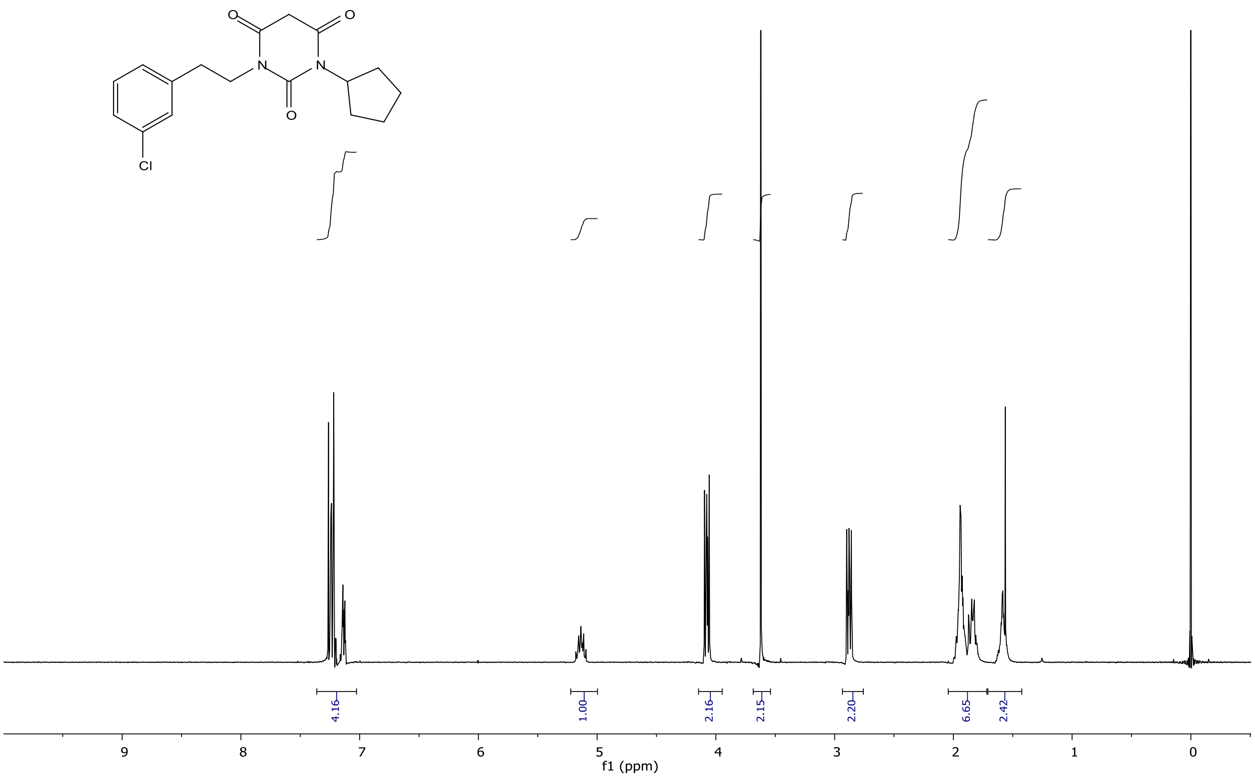


**Figure S10. ¹H NMR** (400 MHz, $CDCl_3$): 1-(3-Chlorophenethyl)-3-cyclopentylpyrimidine-2,4,6-trione **(1)**

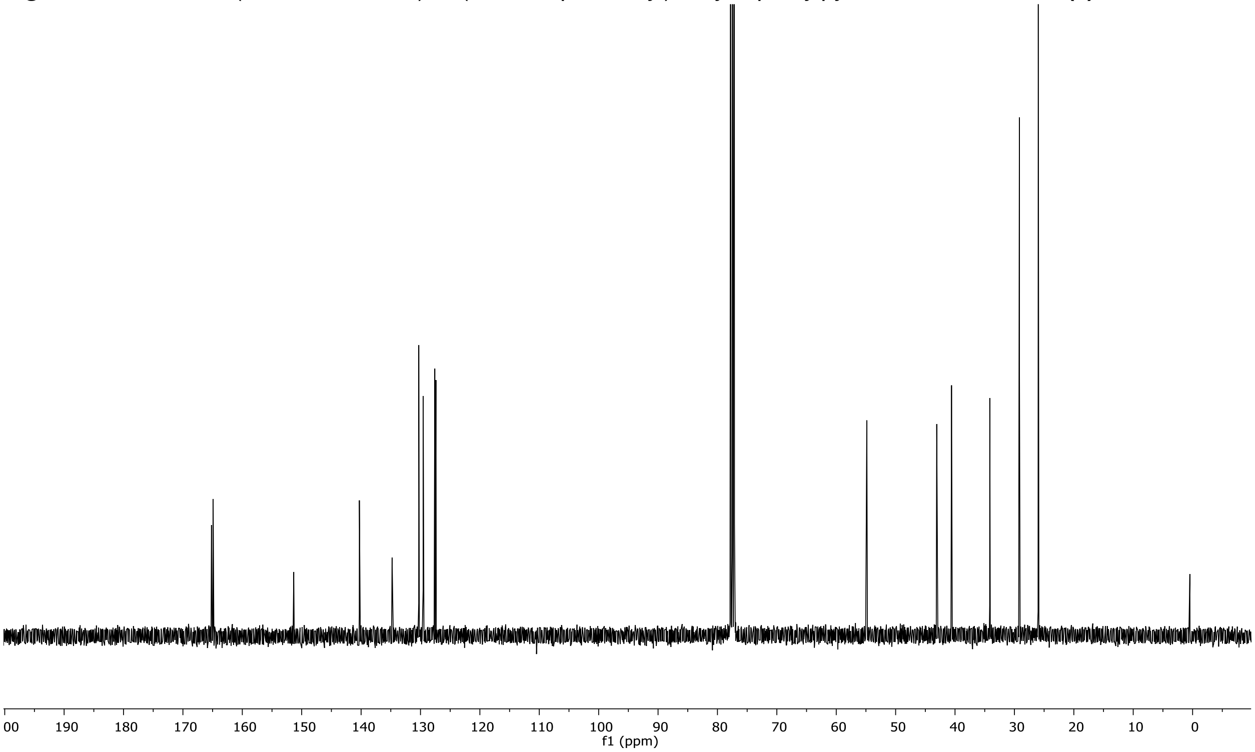


**Figure S11. ¹³C NMR** (101 MHZ, $CDCl_3$): 1-(3-Chlorophenethyl)-3-cyclopentylpyrimidine-2,4,6-trione **(1)**

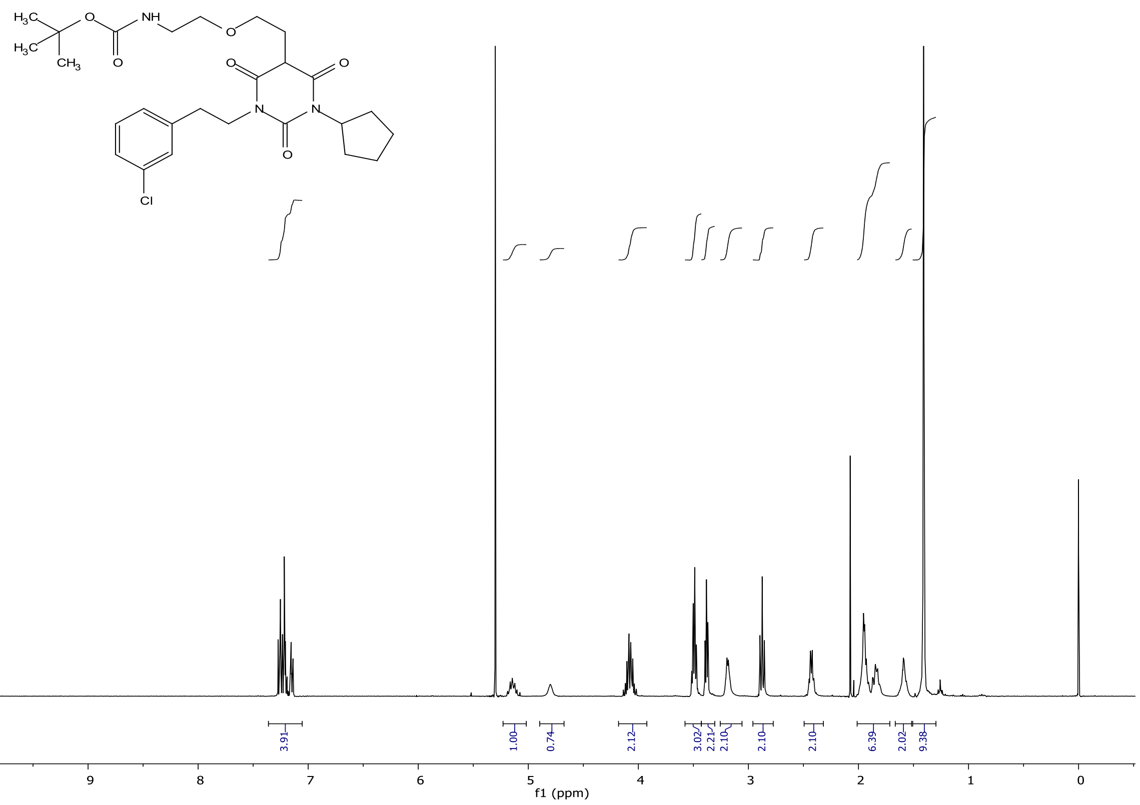


**Figure S12.** $^1$**H NMR** (400 MHz, $CDCl_3$): *tert*-Butyl-(2-(2-(1-cyclopentyl-2,4,6-trioxo-3-phenethylhexahydropyrimidin-5-yl)ethoxy)ethyl)carbamate **(9)**

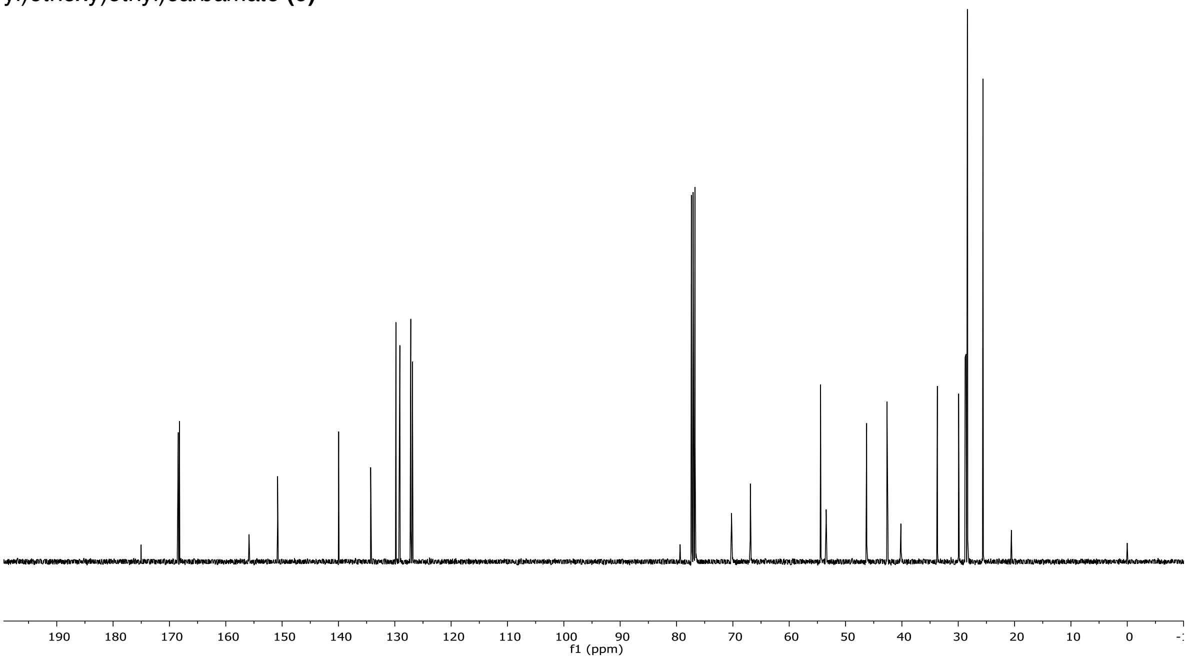


**Figure S13.** $^{13}$**C NMR** (101 MHZ, $CDCl_3$): *tert*-Butyl-(2-(2-(1-cyclopentyl-2,4,6-trioxo-3-phenethylhexahydropyrimidin-5-yl)ethoxy)ethyl)carbamate **(9)**

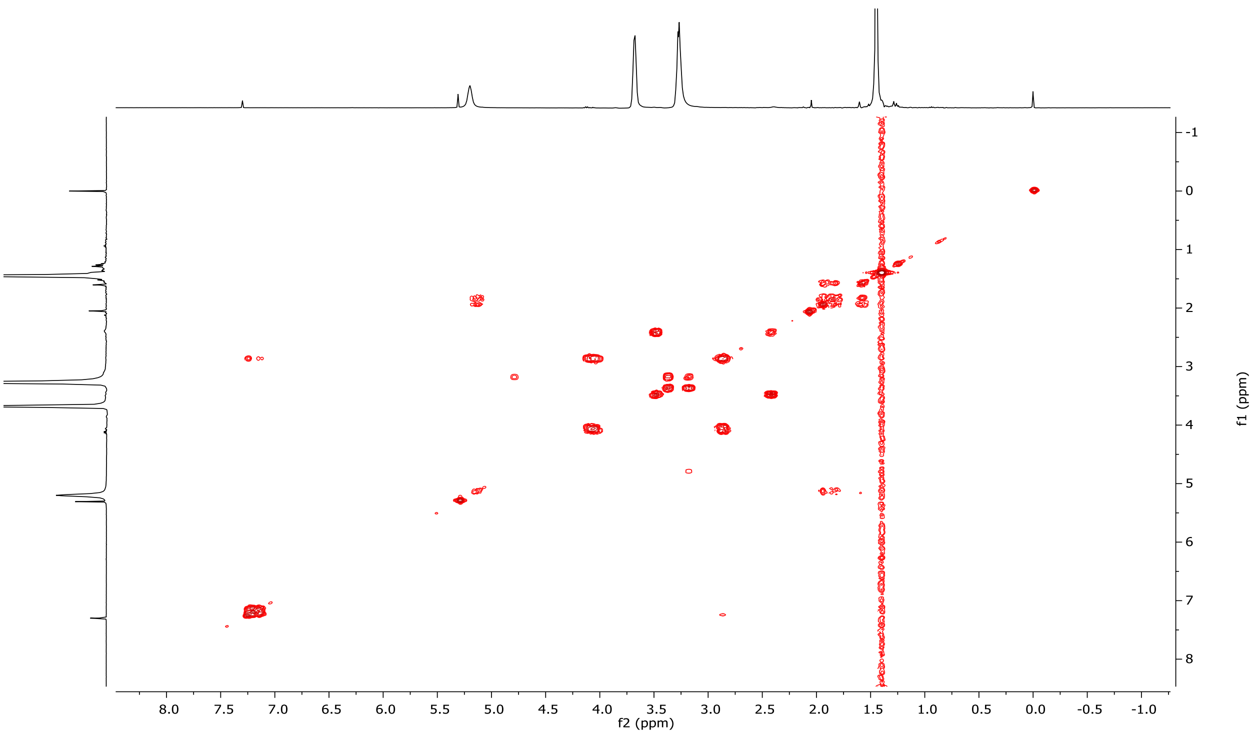


**Figure S14. COSY** (400 MHz, $CDCl_3$): *tert*-Butyl-(2-(2-(1-cyclopentyl-2,4,6-trioxo-3-phenethylhexahydropyrimidin-5-yl)ethoxy)ethyl)carbamate **(9)**

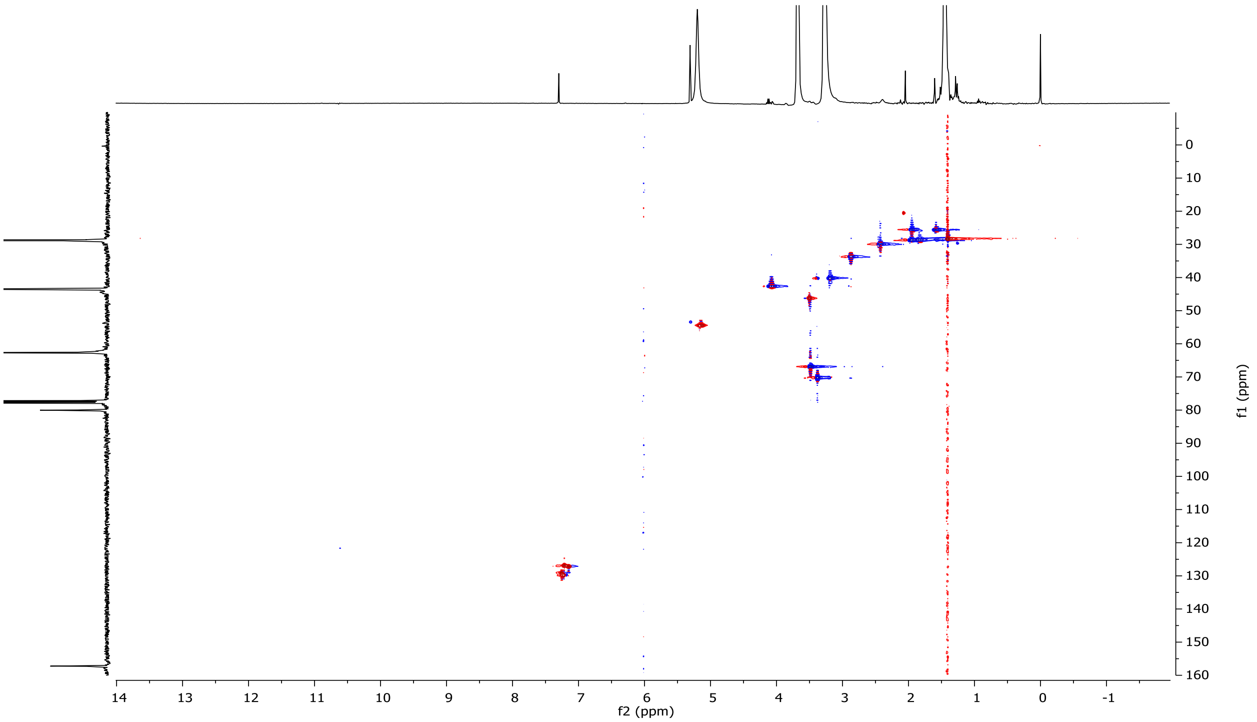


**Figure S15. HSQC** (400 MHz, $CDCl_3$): *tert*-Butyl-(2-(2-(1-cyclopentyl-2,4,6-trioxo-3-phenethylhexahydropyrimidin-5-yl)ethoxy)ethyl)carbamate **(9)**

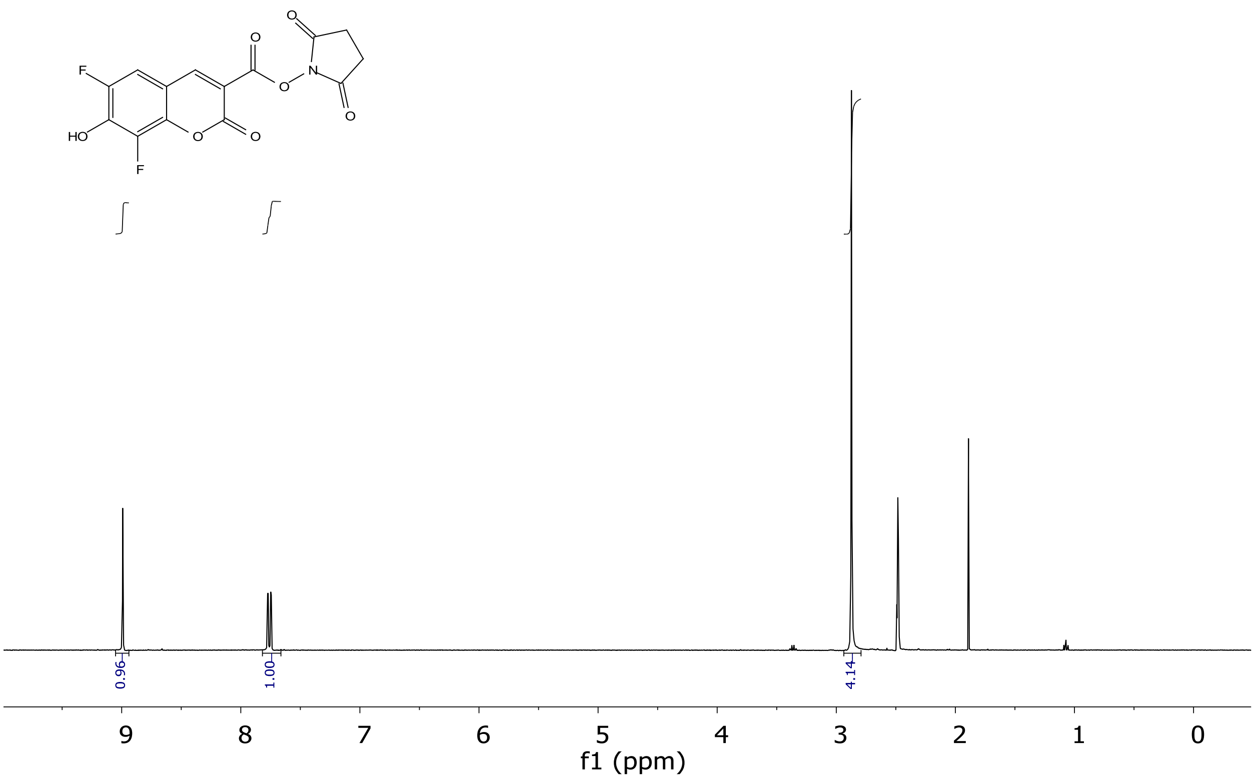


**Figure S16: ¹H NMR** (400 MHz, DMSO-*d₆*)**:** Succinimidyl 6,8-difluoro-7-hydroxycoumarin-3-carboxylate (**11**)

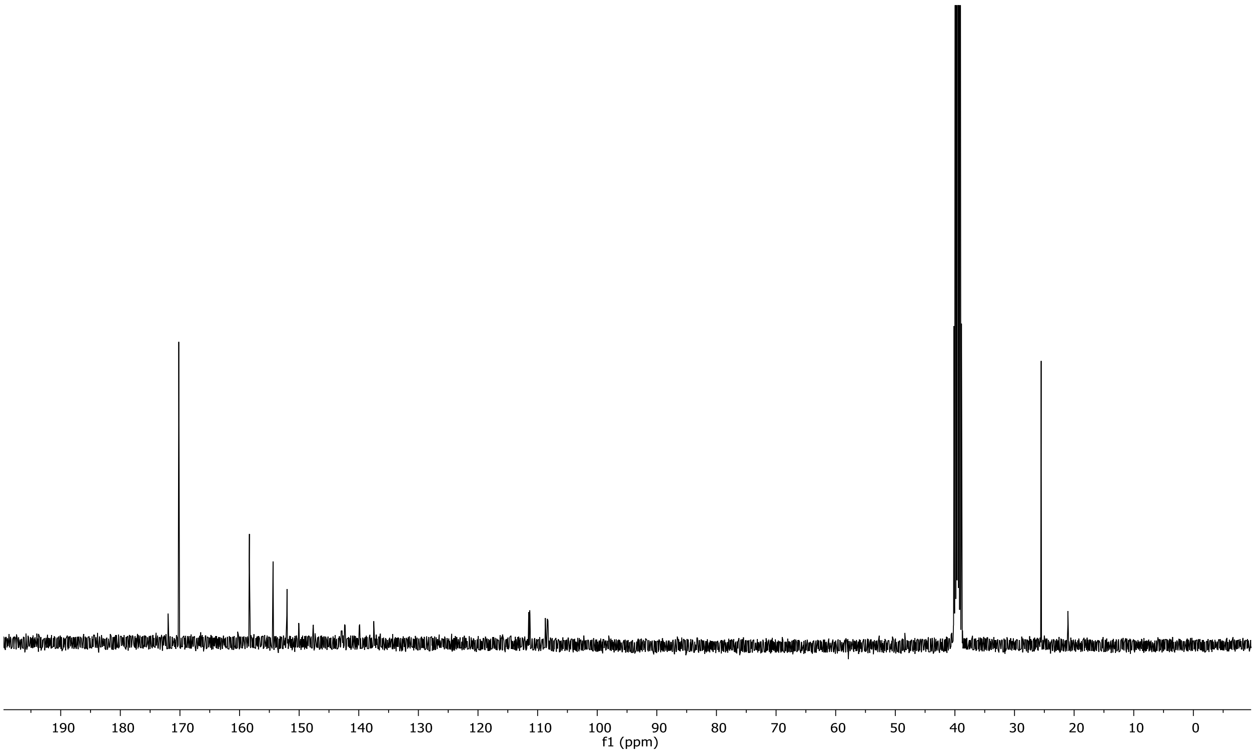


**Figure S17: ¹³C NMR** (101 MHz, DMSO-*d₆*): Succinimidyl 6,8-difluoro-7-hydroxycoumarin-3-carboxylate (**11**)

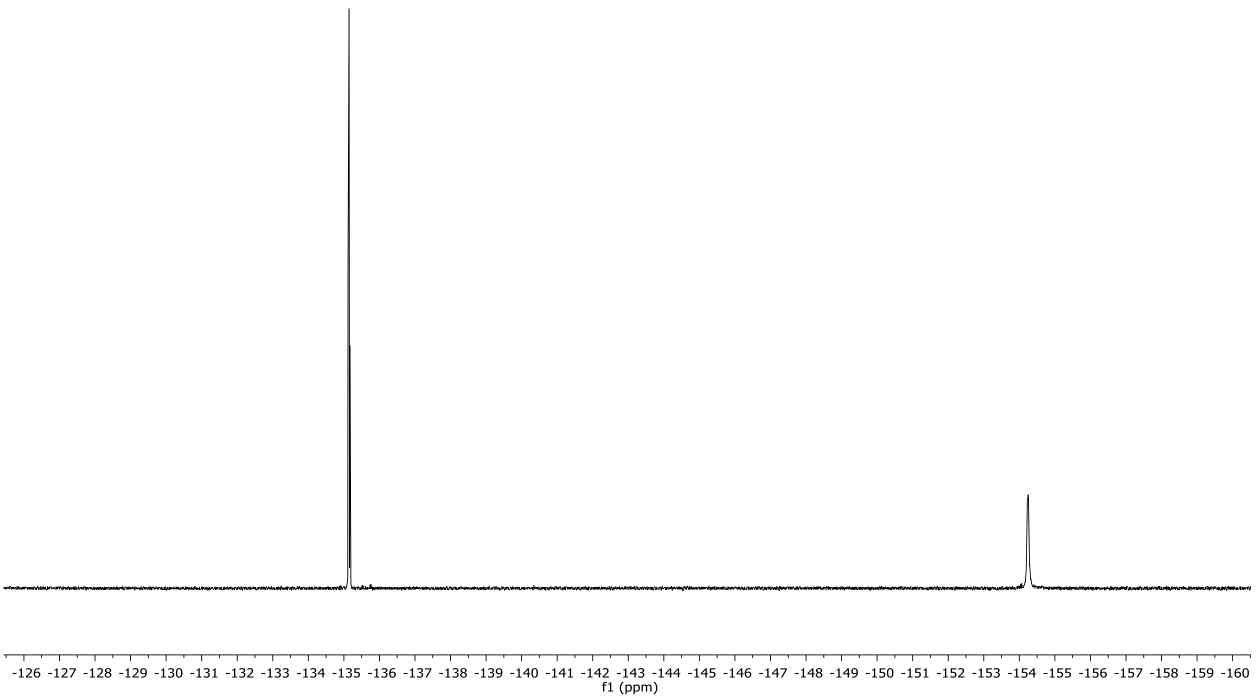


**Figure S18:** $^{19}$**F NMR** (377 MHz, DMSO-$d_6$): Succinimidyl 6,8-difluoro-7-hydroxycoumarin-3-carboxylate (**11**)

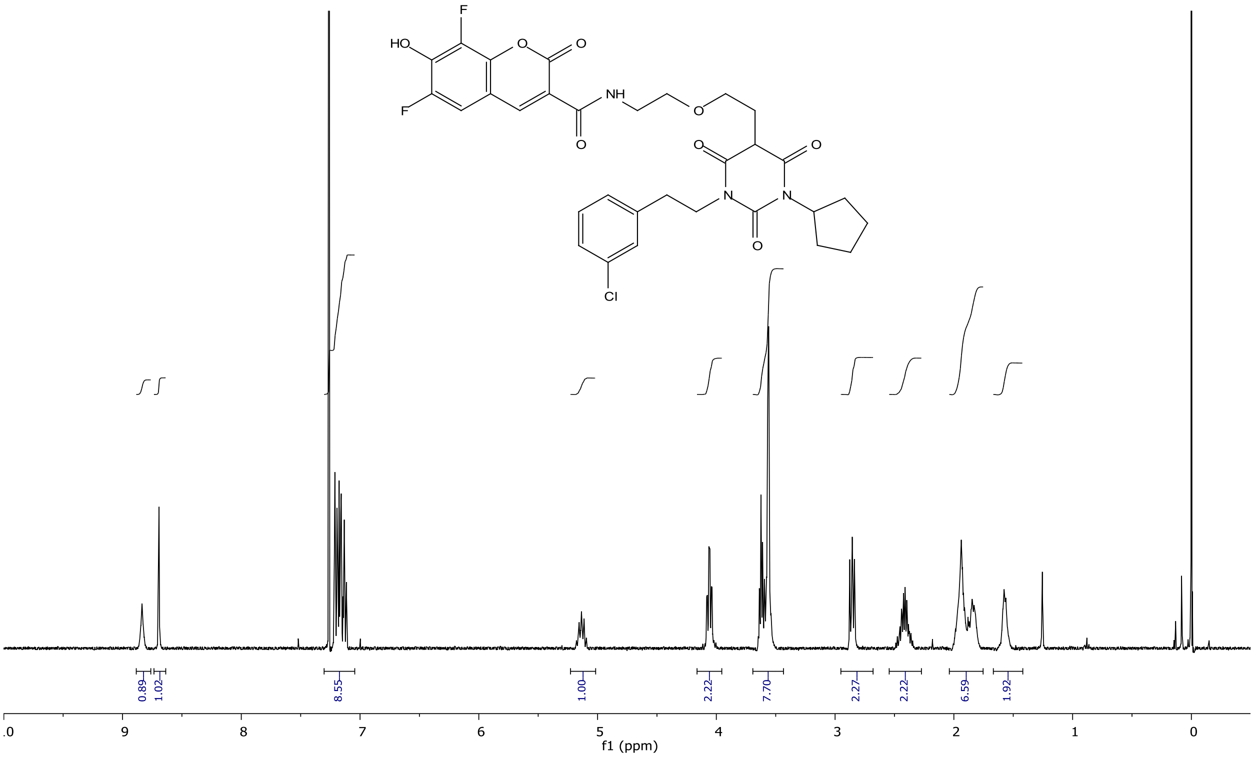


**Figure S19.** $^{1}$**H NMR** (400 MHz, $CDCl_3$): *N*-(2-(2-(1-(3-Chlorophenethyl)-3-cyclopentyl-2,4,6-trioxohexahydropyrimidin-5-l)ethoxy)ethyl)-6,8-difluoro-7-hydroxycoumarin-3-carboxamide **(12)**

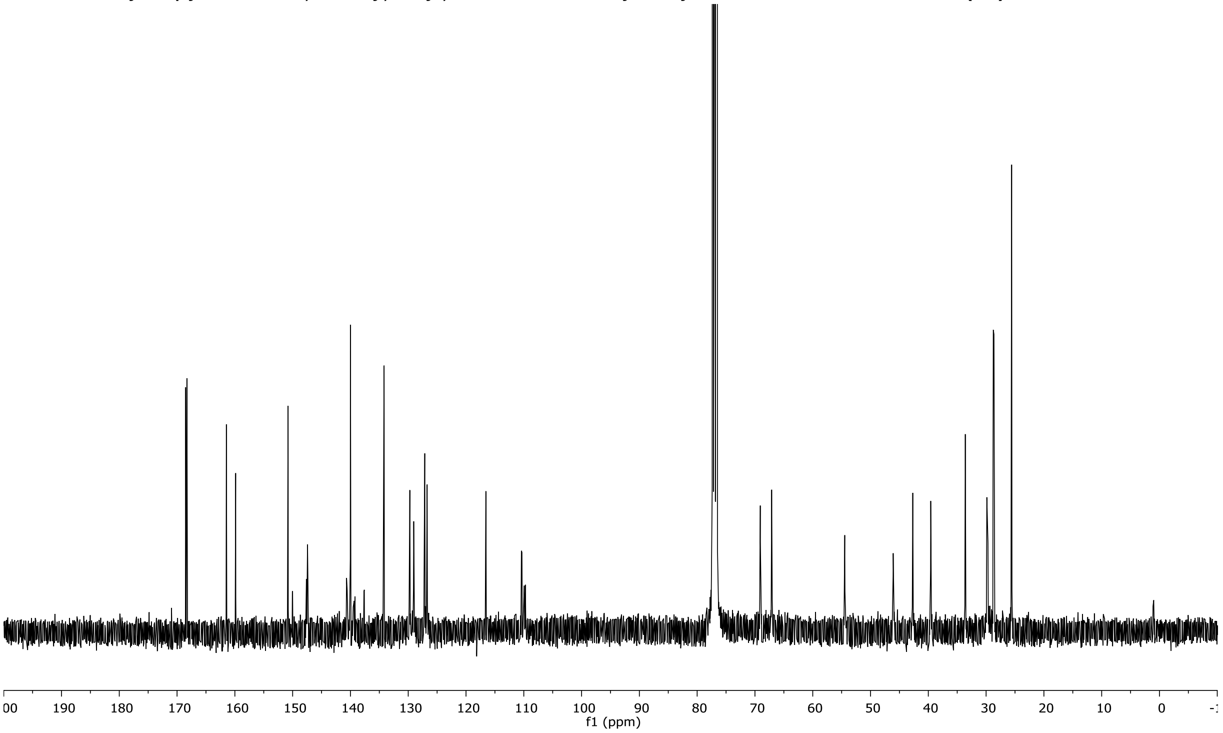


**Figure S20.** $^{13}$**C NMR** (101 MHZ, $CDCl_3$): *N*-(2-(2-(1-(3-Chlorophenethyl)-3-cyclopentyl-2,4,6-trioxohexahydropyrimidin-5-yl)ethoxy)ethyl)-6,8-difluoro-7-hydroxycoumarin-3-carboxamide **(12)**

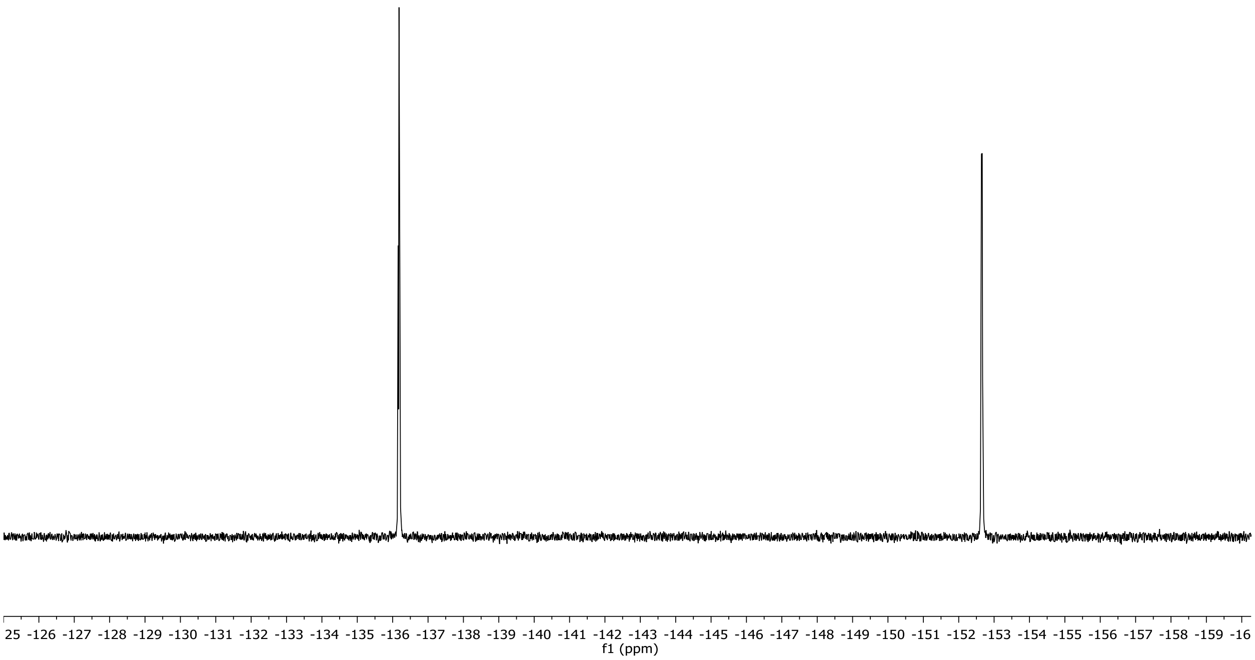


**Figure S21. $^{19}$F NMR** (377 MHz, $CDCl_3$): *N*-(2-(2-(1-(3-Chlorophenethyl)-3-cyclopentyl-2,4,6-trioxohexahydropyrimidin-5-yl)ethoxy)ethyl)-6,8-difluoro-7-hydroxycoumarin-3-carboxamide **(12)**

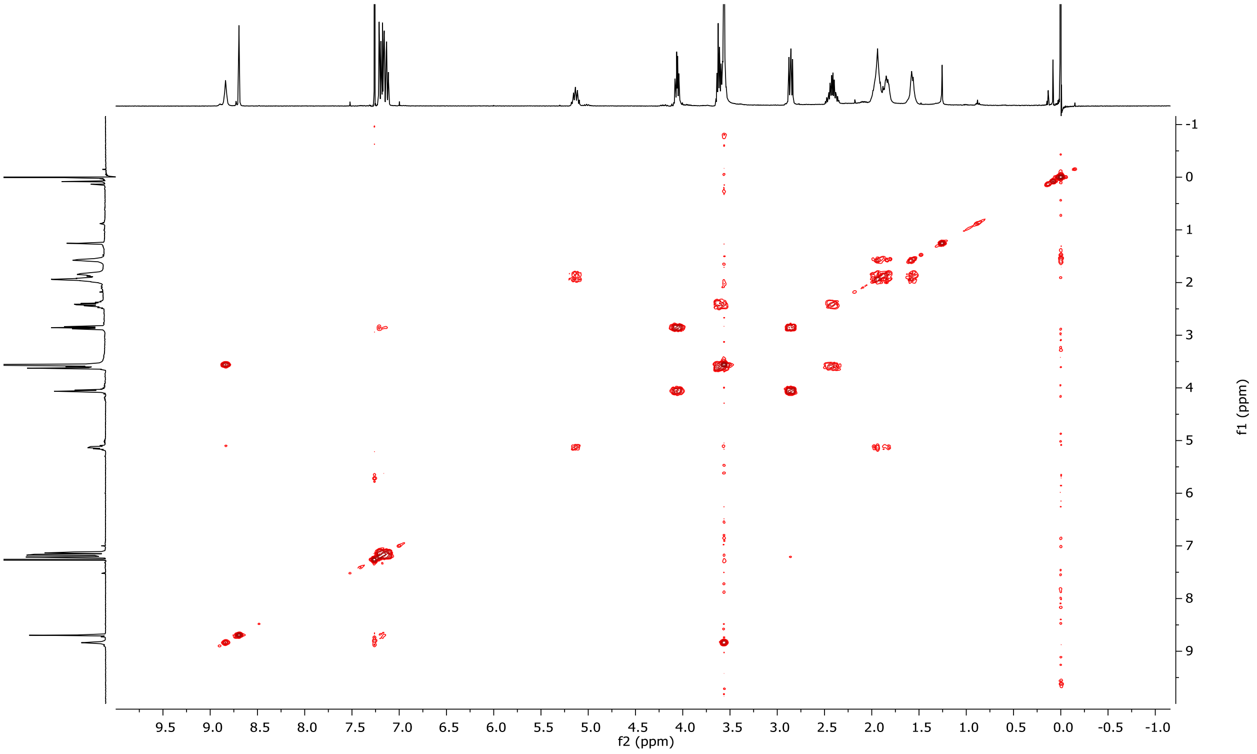


**Figure S22. COSY** (400 MHz, $CDCl_3$): *N*-(2-(2-(1-(3-Chlorophenethyl)-3-cyclopentyl-2,4,6-trioxohexahydropyrimidin-5-yl)ethoxy)ethyl)-6,8-difluoro-7-hydroxycoumarin-3-carboxamide **(12)**

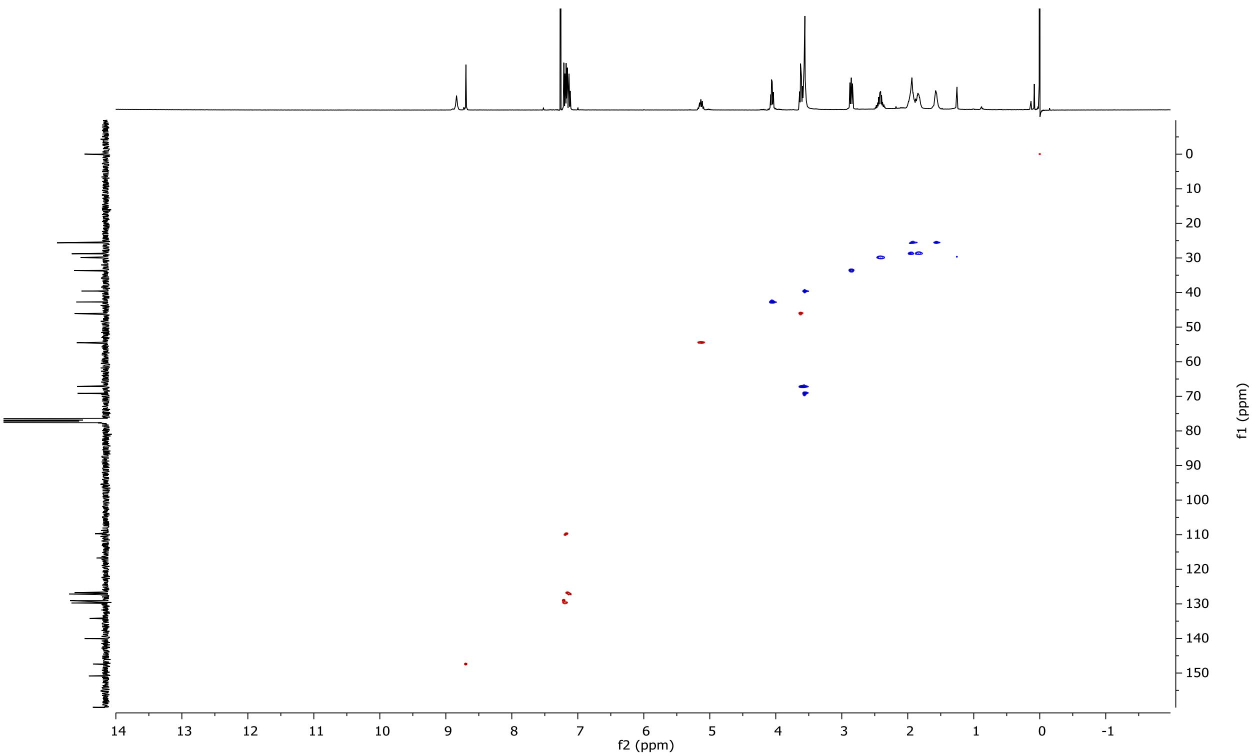


**Figure S23. HSQC** (400 MHz, $CDCl_3$): *N*-(2-(2-(1-(3-Chlorophenethyl)-3-cyclopentyl-2,4,6-trioxohexahydropyrimidin-5-yl)ethoxy)ethyl)-6,8-difluoro-7-hydroxycoumarin-3-carboxamide **(12)**
